# Males are more sensitive to their audience than females when scent-marking in the redfronted lemur

**DOI:** 10.1101/2021.09.02.458772

**Authors:** Louise R. Peckre, Alexandra Michiels, Lluís Socias-Martínez, Peter M. Kappeler, Claudia Fichtel

## Abstract

Audience effects, i.e. changes in behaviour caused by the presence of conspecifics, have rarely been studied in the context of olfactory communication, even though they may provide important insights into the functions of olfactory signals. Functional sex differences in scent-marking behaviours are common and influenced by the social system. To date, patterns of functional sex differences in scent-marking behaviours remain unknown in species without overt dominance relationships. We investigated sex differences in intra-group audience effects on anogenital scent-marking in a wild population of redfronted lemurs (*Eulemur rufifrons*) by performing focal scent-marking observations. With a combination of generalised linear mixed models and exponential random graph models, we found different audience effects in both sexes. Males were overall more sensitive than females to their audience. Only males seemed to be sensitive to the presence of both members of the opposite sex and same-sex conspecifics in the audience. Females were only moderately sensitive to the presence of other females in the audience. This study offers a potential behavioural pattern associated with anogenital scent-marking that seem to differ from those described for species exhibiting female dominance, supporting the notion that the social systems co-varies with scent-marking behaviours and scent-complexity in strepsirrhines.

## Introduction

The traditional approach of considering communication as information transfer between a signaller-receiver dyad connected by a transmission channel^1^ has been extended by the concept of communication networks. Indeed, in many social groups, individuals are closely spaced, signals reaching multiple individuals, including both intended and unintended receivers^2–4^. Unintended receivers, i.e. eavesdroppers, can exploit information to their benefit, sometimes at a cost to the signaller^3,5^. Accordingly, signallers may be sensitive to the presence and characteristics of receivers and may exhibit behavioural flexibility by initiating, inhibiting, or varying the rate or nature of signal production. Such ‘audience effects’ have mainly been described for vocal and visual signals^6^. In contrast, audience effects on the transmission of olfactory signals remain poorly studied, even though scent represents the main modality of communication in most mammals^5,7^.

Scent-marking behaviours are defined as specialised motor patterns used to deposit chemical secretions or excretion (e.g., urine, saliva, anogenital secretions) on objects or conspecifics^8–10^. These signals can carry reliable information of the sender’s age, sex, health, reproductive and social status^11–15^. Scent marking behaviours have been associated with various functions, both across^16–19^ and within species^20–23^. They can be classified into three broad functional categories: sexual attraction, competition, and parental care^16–19,24^. For instance, in rodents, female secretions play a role in coordinating reproduction with males, suppressing reproduction in other females, territory advertisement and defence, as well as in maternal care^18^.

Audience effects in olfactory signals may have been rarely studied because scent signals, in contrast to visual and acoustic signals, are long-lasting, remaining in the environment long after the sender left the location. Hence, signals are exposed to an audience present during scent deposition but also to an unknown audience long after deposition, resulting in some uncertainty about the identity of the receivers. However, scent-marks are often associated with conspicuous ephemeral visual displays when deposited. These visual components might immediately attract the attention of individuals present in the vicinity and guide them to the signal’s olfactory component, allowing senders some control over the identity of receivers. Hence, this multimodal nature may confer scent-marking behaviour the capacity to be addressed both to the audience present during deposition and an unknown future audience^5,20,24^. This idea was conceptually formalised twice. First, the ‘demonstrative marking hypothesis’ was postulated in territorial male Thomson’s gazelles (*Eudorcas thomsonii*), which associate urine-faeces deposition with an extreme body posture display^25–27^. Later, the same idea has been described as a ‘composite effect’ in ring-tailed lemurs (*Lemur catta*), which either urinate with the tail only slightly raised or combine urine-marking with a conspicuous visual signal, the up-right erection of the tail. The erection of their tail attracted the visual attention of receivers to the location of the urine deposition and resulted in more group members inspecting the urine-mark, compared to urine-marks deposited without tail display^28^.

Testing for potential audience effects on scent-marking behaviour might also be a promising approach to explore the functions and potential sex differences associated with scent-marking behaviours. Functional differences in scent-marking behaviours between sexes have been described in several species (e.g. mandrills^29^, cheetahs^23^). These functional differences are usually associated with morphological, physiological and behavioural differences. Sex differences in the glandular structure and the frequency of scent-marking behaviours are indeed common^13,19,20,23,30–38^. The balance of investment in chemical signalling and scents’ deployment across sexes seems to be influenced by the species’ social system. For instance, male-biased dimorphism is often reduced in monogamous species, and female-biased dimorphism occurs in species with high levels of female competition^16,19,39–41^. However, only very few studies investigated how a species’ social system influences scent-marking behaviours and functionality^42–44^.

Strepsirrhines primates, like most other mammals (but unlike anthropoid primates), have a functional vomeronasal organ. They rely heavily on olfactory communication and produce a wide variety of chemical signals expressed by glands located in various body areas (head, neck, chest, forelimb and anogenital area)^24,36,45,46^. Interestingly, diversification of means of olfactory communication in strepsirrhines evolved along with broad diversification of their ecology and social systems^46^.

The genus *Eulemur* (Lemuridae) includes five species exhibiting female dominance, a common pattern in strepsirrhines, and four species without overt dominance relations between sexes^47–49^ (sometimes referred to as codominant species). Species exhibiting female dominance evolved more elaborate anogenital glands (often more elaborate than those of males), produce scents chemically more complex than those of males and exhibit more pronounced scent-marking behaviour associated with a dominance status signalling function^41,48^ as also found in other lemur species exhibiting female dominance^14,22,50^. Species in which none of the sex is overtly dominant over the other exhibit conspicuous anogenital scent-marking displays with no sexual bias in the frequency of anogenital scent-marking^41,51^. However, even though females seem to have more elaborate glandular folds, the chemical richness of perianal and genital secretions in these species is higher in males than in females^36,48^. These morphological and physiological differences may be indicative of functional differences. However, the pattern of functional sex differences remains unstudied in the species without overt intersexual dominance relations. Moreover, these species also show low aggression levels, without clear dominance patterns within sexes neither, and are, as such, qualified as egalitarian, which compromises the existence of a status signalling function associated with a dominance status in these species.

In this study, we investigated anogenital scent-marking behaviour in wild redfronted lemurs (*E. rufifrons*). Redfronted lemurs live in cohesive small multi-female–multi-male groups of 5-12 individuals with an even or male-biased sex ratio^52–56^. They lack a clear pattern of intersexual dominance relationships, with none of the sexes being consistently dominant over the other^57,58^ and aggression rates are low within both sexes^59^. We examined whether males and females differed in their sensitivity to an audience present in a 3, 5, and 10 m radius when scent-marking and whether these potential differences may reveal functional sex differences associated with anogenital scent-marking in this species.

On the one hand, at the intersexual level, scent-marking signals have first been suggested to serve a pair-bonding maintenance function, as shown in both pair-living (*Eulemur rubriventer*^60^) and group-living species (*Propithecus coquereli*^61^)(Table 1). Red-fronted lemurs being promiscuous, with all females mating with virtually all males within their group^62^ and no strong male-female bonds being observed^58^, this pair-bonding maintenance function of scent-marking is unlikely. However, scent-marking signals have also been suggested to be directed to the opposite sex as a form of mate attraction (Table 1). If a form of mate attraction occurs, we would expect one sex to increase its probability to scent-mark in the presence of an increased proportion of individuals of the opposite sex. In redfronted lemurs, one central male in each group seems to be mainly associated with all females^58^. Central males sire around 60-70% of all infants^54,63^ and scent-mark more than other males in the group. Hence, we predict (I) that males will perform more anogenital scent-marking in the presence of females and that this effect should be dependent on the social relationships the male maintains with the females present in the audience.

**Table 1:** Functions suggested for anogenital scent-marking behaviours in strepsirrhines depending on the social structure and the sex of individuals.

|  |  | Female dominance & hierarchy in both sexes |  | No dominance relationships both between and within sexes |  |
| --- | --- | --- | --- | --- | --- |
| Function |  | Females | Males | Females | Males |
| Sexual attraction | Advertisement of reproductive receptivity | *Lemur catta*<sup>64</sup> | / | / | / |
|  | Signals of individual quality | / | *Propithecus Verreauxi*<sup>65</sup>, *L. catta*<sup>28</sup>, *Microcebus murinus*<sup>66</sup> | / | / |
|  | Pair-bonding maintenance | *P. coquereli*<sup>61</sup> | *P. coquereli*<sup>61</sup> | / | / |
| Competition | Social status signalling | *P. edwardsi*<sup>35</sup> | *P. candidus*<sup>67</sup>; *P. edwardsi*<sup>35,67</sup>; *P. verreauxi*<sup>22,68</sup>; *L. catta*<sup>69</sup> | *Eulemur Fulvus* & *E. macaco*<sup>51</sup> | *E. fulvus* & *E. macaco*<sup>51</sup>; *E. rufifrons*<sup>58</sup>; *E. rufus*<sup>33</sup> |
|  | Reproductive suppression | *M. murinus*<sup>70</sup> | *M. murinus*<sup>71</sup>; *P. verreauxi*<sup>72</sup> | / | / |
|  | Reproductive synchronisation | *L. catta*<sup>73</sup> | / | / | / |

On the other hand, scent-marking behaviours have been proposed to serve in intra-sexual communication. Individuals may use anogenital scent-marking for advertising their social status as a form of indirect competition (Table 1). Indeed, scent-marking behaviours have more broadly been suggested to be non-overt agonistic status indicators and are, as such, expected to be more prominent in species with low aggression rates^13,74^. In male redfronted lemurs, there is no linear dominance hierarchy, and aggression rates are relatively low, but males still differ in their social value (time spent socialising)^58^. Hence, we predict (II) that male redfronted lemurs should modify their scent-marking behaviour in response to a male audience and that this effect should be particularly dependent on their social value and the one of the males in the audience. Female redfronted lemurs also do not exhibit hierarchical dominance relationships. However, competition can be intense, with females evicting even related females from the group when they reach a critical group size of about ten individuals^75^. Hence, similarly to what we predicted for males, we predict (III) that females should modify their scent-marking behaviour in response to a female audience and that this effect should be particularly dependent on their social value and the one of the females in the audience.

## Results

In total, we recorded a total of 327 events on scent-marking spots corresponding to 223 anogenital scent-marking events (105 in males and 118 in females) and 104 pass by events (i.e. without scent-marking)(60 in males and 44 in females).

In males, we found a significant audience effect within the 3 m radius (full-null model comparison: χ^2^=6.48, df=2, P=0.039; R^2^_m_=0.10, R^2^_c_=0.23). Notably, males anogenital marked less often when an increased proportion of males were present (χ^2^=6.23, df=1, P=0.013, Table 2, Fig. 1a). However, this audience effect was not observed within the 5 and 10 m radius (full-null model comparisons: for 5 m, χ^2^=4.77, df=2, P=0.092, Fig. 1b; for 10 m, χ^2^=1.47, df=2, P=0.481, Table 2, Fig. 1c). In addition, for all three distance radiuses, neither the proportion of females, age, context, nor season significantly affected the probability of anogenital marking in males (Table 2).

**Table 2:**
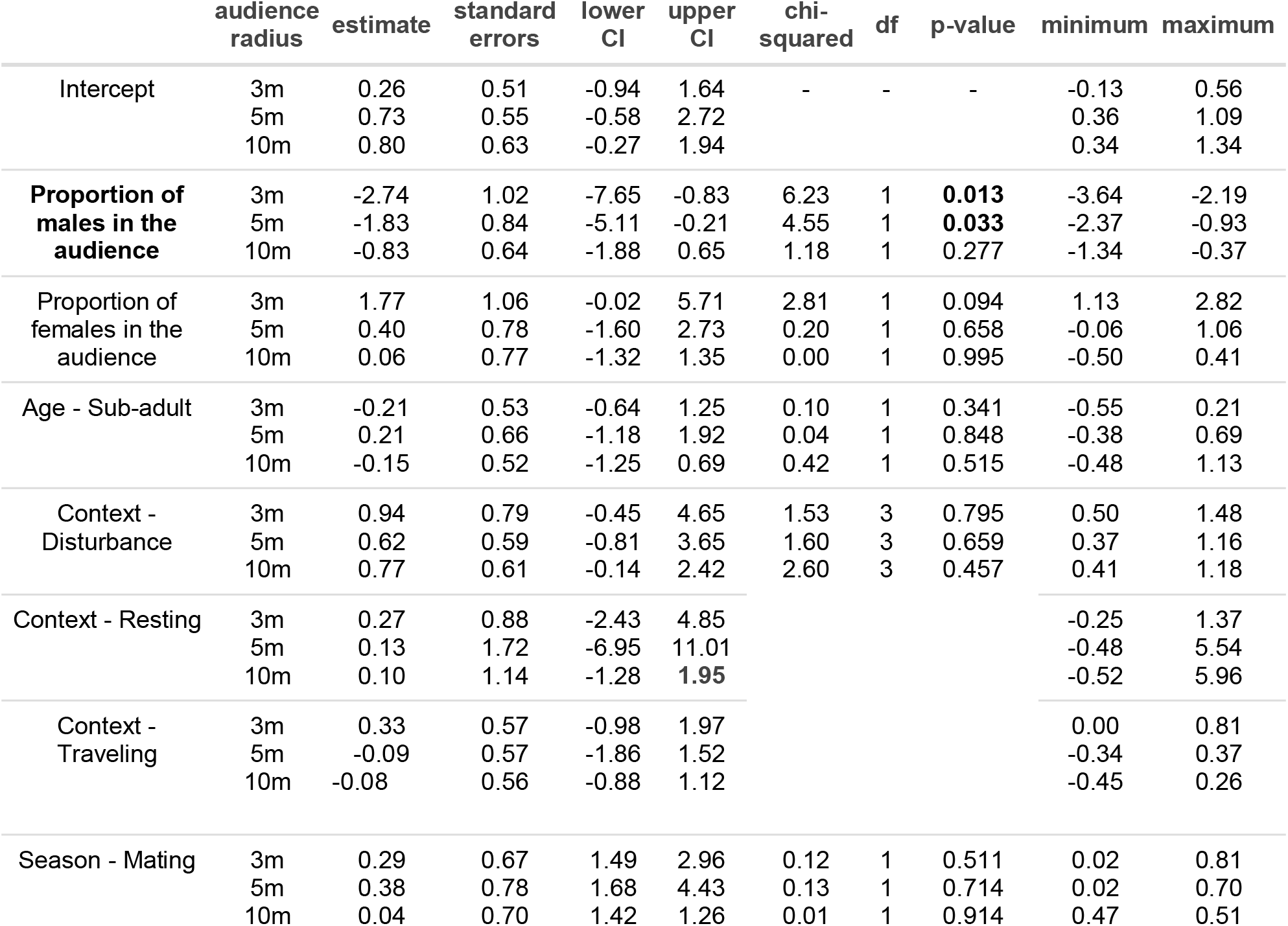
Results of the models of the effects of audience composition within 3, 5 and 10m radius, age, context and season on the probability that a male scent-mark when passing a marking spot.

|  | audience radius | estimate | standard errors | lower CI | upper CI | chi-squared | df | p-value | minimum | maximum |
| --- | --- | --- | --- | --- | --- | --- | --- | --- | --- | --- |
| Intercept | 3m | 0.26 | 0.51 | -0.94 | 1.64 | - | - | - | -0.13 | 0.56 |
|  | 5m | 0.73 | 0.55 | -0.58 | 2.72 |  |  |  | 0.36 | 1.09 |
|  | 10m | 0.80 | 0.63 | -0.27 | 1.94 |  |  |  | 0.34 | 1.34 |
| **Proportion of males in the audience** | 3m | -2.74 | 1.02 | -7.65 | -0.83 | 6.23 | 1 | **0.013** | -3.64 | -2.19 |
|  | 5m | -1.83 | 0.84 | -5.11 | -0.21 | 4.55 | 1 | **0.033** | -2.37 | -0.93 |
|  | 10m | -0.83 | 0.64 | -1.88 | 0.65 | 1.18 | 1 | 0.277 | -1.34 | -0.37 |
| Proportion of females in the audience | 3m | 1.77 | 1.06 | -0.02 | 5.71 | 2.81 | 1 | 0.094 | 1.13 | 2.82 |
|  | 5m | 0.40 | 0.78 | -1.60 | 2.73 | 0.20 | 1 | 0.658 | -0.06 | 1.06 |
|  | 10m | 0.06 | 0.77 | -1.32 | 1.35 | 0.00 | 1 | 0.995 | -0.50 | 0.41 |
| Age - Sub-adult | 3m | -0.21 | 0.53 | -0.64 | 1.25 | 0.10 | 1 | 0.341 | -0.55 | 0.21 |
|  | 5m | 0.21 | 0.66 | -1.18 | 1.92 | 0.04 | 1 | 0.848 | -0.38 | 0.69 |
|  | 10m | -0.15 | 0.52 | -1.25 | 0.69 | 0.42 | 1 | 0.515 | -0.48 | 1.13 |
| Context - Disturbance | 3m | 0.94 | 0.79 | -0.45 | 4.65 | 1.53 | 3 | 0.795 | 0.50 | 1.48 |
|  | 5m | 0.62 | 0.59 | -0.81 | 3.65 | 1.60 | 3 | 0.659 | 0.37 | 1.16 |
|  | 10m | 0.77 | 0.61 | -0.14 | 2.42 | 2.60 | 3 | 0.457 | 0.41 | 1.18 |
| Context - Resting | 3m | 0.27 | 0.88 | -2.43 | 4.85 |  |  |  | -0.25 | 1.37 |
|  | 5m | 0.13 | 1.72 | -6.95 | 11.01 |  |  |  | -0.48 | 5.54 |
|  | 10m | 0.10 | 1.14 | -1.28 | 1.95 |  |  |  | -0.52 | 5.96 |
| Context - Traveling | 3m | 0.33 | 0.57 | -0.98 | 1.97 |  |  |  | 0.00 | 0.81 |
|  | 5m | -0.09 | 0.57 | -1.86 | 1.52 |  |  |  | -0.34 | 0.37 |
|  | 10m | -0.08 | 0.56 | -0.88 | 1.12 |  |  |  | -0.45 | 0.26 |
| Season - Mating | 3m | 0.29 | 0.67 | 1.49 | 2.96 | 0.12 | 1 | 0.511 | 0.02 | 0.81 |
|  | 5m | 0.38 | 0.78 | 1.68 | 4.43 | 0.13 | 1 | 0.714 | 0.02 | 0.70 |
|  | 10m | 0.04 | 0.70 | 1.42 | 1.26 | 0.01 | 1 | 0.914 | 0.47 | 0.51 |

**Figure 1:**
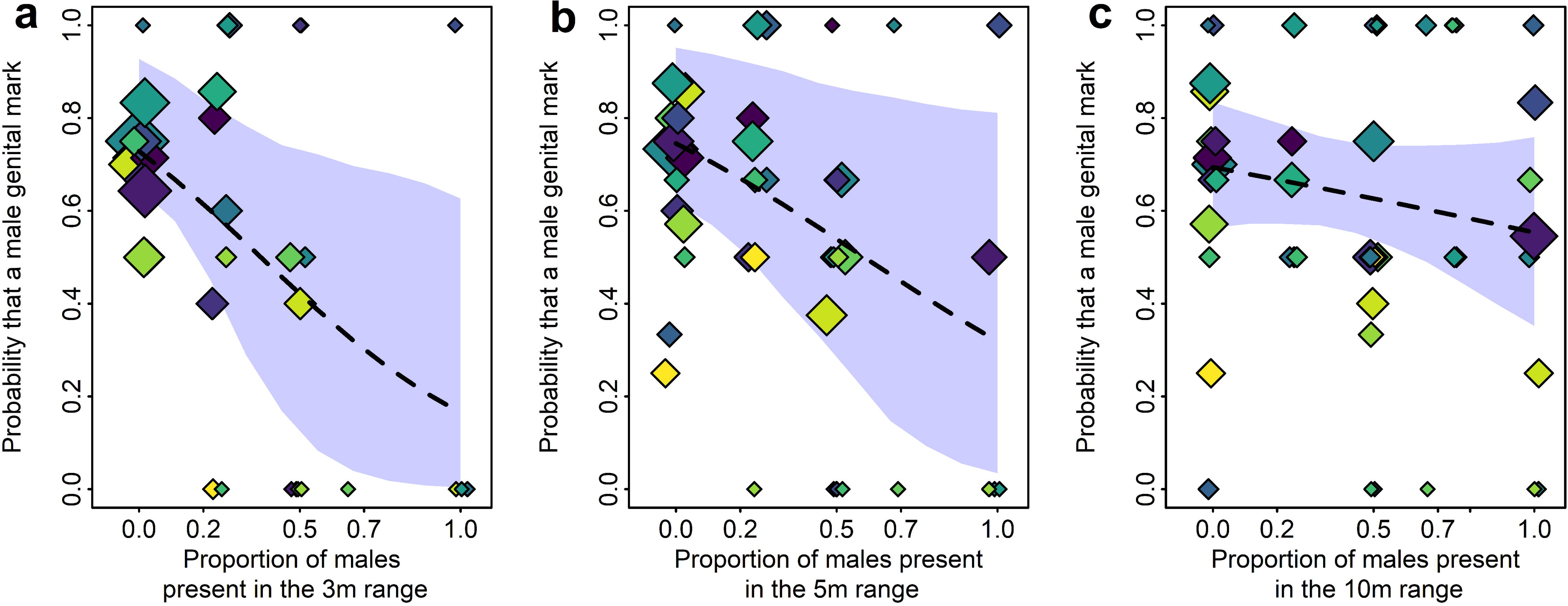
Probability that a male deposited a scent-mark depending on the proportion of males present a) in a 3m radius, b) in a 5m radius, c) in a 10m radius. Colours correspond to the different individuals (n=14), and the size of the circle corresponds to the number of observations (in total, n=165).

In females, we found no audience effect associated with the proportion of individuals present observed (full-null model comparison: for 3m, χ^2^=4.84, df=2, P=0.089; for 5 m, χ^2^=6.85, df=2, P=0.032; for 10m χ^2^=3.54, df=2, P=0.171; Table 3). Neither the proportion of males, the proportion of females, age, context, nor season predicted the probability of anogenital scent-marking (Table 3).

**Table 3:** Results of the models of the effects of audience composition within 3, 5 and 10m radius, age, context and season on the probability that a female scent-marked when passing a marking spot.

|  | audience radius | estimate | standard errors | lower CI | upper CI | chi-squared | df | p-value | minimum | maximum |
| --- | --- | --- | --- | --- | --- | --- | --- | --- | --- | --- |
| Intercept | 3m | 1.41 | 0.39 | 0.71 | 2.44 | - | - | - | -0.17 | 0.48 |
|  | 5m | 1.61 | 0.45 | 0.89 | 2.99 |  |  |  | 1.46 | 2.00 |
|  | 10m | 1.95 | 0.68 | 0.99 | 5.00 |  |  |  | 1.68 | 2.40 |
| Proportion of males in the audience | 3m | -0.67 | 0.84 | -2.62 | 1.34 | 0.64 | 1 | 0.425 | -3.67 | -2.12 |
|  | 5m | -0.50 | 0.85 | -2.49 | 1.35 | 0.34 | 1 | 0.558 | -1.05 | -0.07 |
|  | 10m | -1.75 | 1.00 | -5.45 | -0.06 | 3.01 | 1 | 0.083 | -2.67 | -1.34 |
| Proportion of females in the audience | 3m | -0.71 | 0.71 | -2.50 | 0.78 | 1.00 | 1 | 0.319 | 0.95 | 2.79 |
|  | 5m | -0.95 | 0.66 | -2.60 | 0.48 | 2.13 | 1 | 0.145 | -1.29 | -0.55 |
|  | 10m | 0.23 | 0.73 | -1.48 | 2.45 | 0.11 | 1 | 0.745 | 0.03 | 0.96 |
| Context - Disturbance | 3m | -0.10 | 0.53 | -1.20 | 1.23 | 1.01 | 3 | 0.799 | 0.11 | 0.76 |
|  | 5m | -0.09 | 0.53 | -1.24 | 1.27 | 1.30 | 3 | 0.730 | -0.44 | 0.17 |
|  | 10m | -0.07 | 0.56 | -1.73 | 2.05 | 1.71 | 3 | 0.634 | -0.35 | 0.30 |
| Context - Resting | 3m | 0.34 | 0.64 | -0.92 | 2.75 |  |  |  | 0.35 | 1.36 |
|  | 5m | 0.18 | 0.65 | -1.30 | 2.53 |  |  |  | -0.14 | 0.92 |
|  | 10m | 0.55 | 0.79 | -1.11 | 6.57 |  |  |  | 0.21 | 1.44 |
| Context - Traveling | 3m | -0.30 | 0.49 | -1.44 | 0.80 |  |  |  | -0.37 | 1.45 |
|  | 5m | -0.47 | 0.51 | -1.79 | 0.64 |  |  |  | -0.81 | -0.26 |
|  | 10m | -0.55 | 0.61 | -2.65 | 0.91 |  |  |  | -0.92 | -0.29 |
| Season - Mating | 3m | -0.13 | 0.48 | -1.18 | 1.29 | 0.07 | 1 | 0.790 | -0.03 | 0.73 |
|  | 5m | -0.02 | 0.49 | -1.08 | 1.35 | 0.00 | 1 | 0.962 | -0.29 | 0.22 |
|  | 10m | -0.14 | 0.59 | -1.69 | 2.21 | 0.06 | 1 | 0.813 | -0.38 | 0.16 |

When considering the anogenital-marking network (exponential random graph model), overall, sex and/or sociality (i.e. DSI rank of the dyad focal-audience, difference in the CSI values of the individuals within a given dyad, CSI of the individual in the audience) had an effect on the probability of an individual to scent-mark in front of another individual (full-null model comparison: 3m chi2=-697.5, df=580, p<0.001; 5m chi2=-1263.1, df=580, p<0.001; 10m chi2=-1982.8, df=580, p<0.001).

In particular, there was a significant effect of the interaction between the combination of sexes and the DSI rank of the respective dyad on the probability of scent-marking within the 5m and 10m radius (full-reduced model comparison: 5m chi2=-1412.5, df=568, p-value=0.003; 10m chi2=-2213.5, df=568, p-value<0.001; Table 4) but not within the 3m radius (chi2=-749.5, df=568, p-value=0.184). Males scent-marked more often in front of females with whom they interacted more often. In contrast, females scent-marked more often in front of females with whom they interacted less (Fig. 2).

**Table 4:**
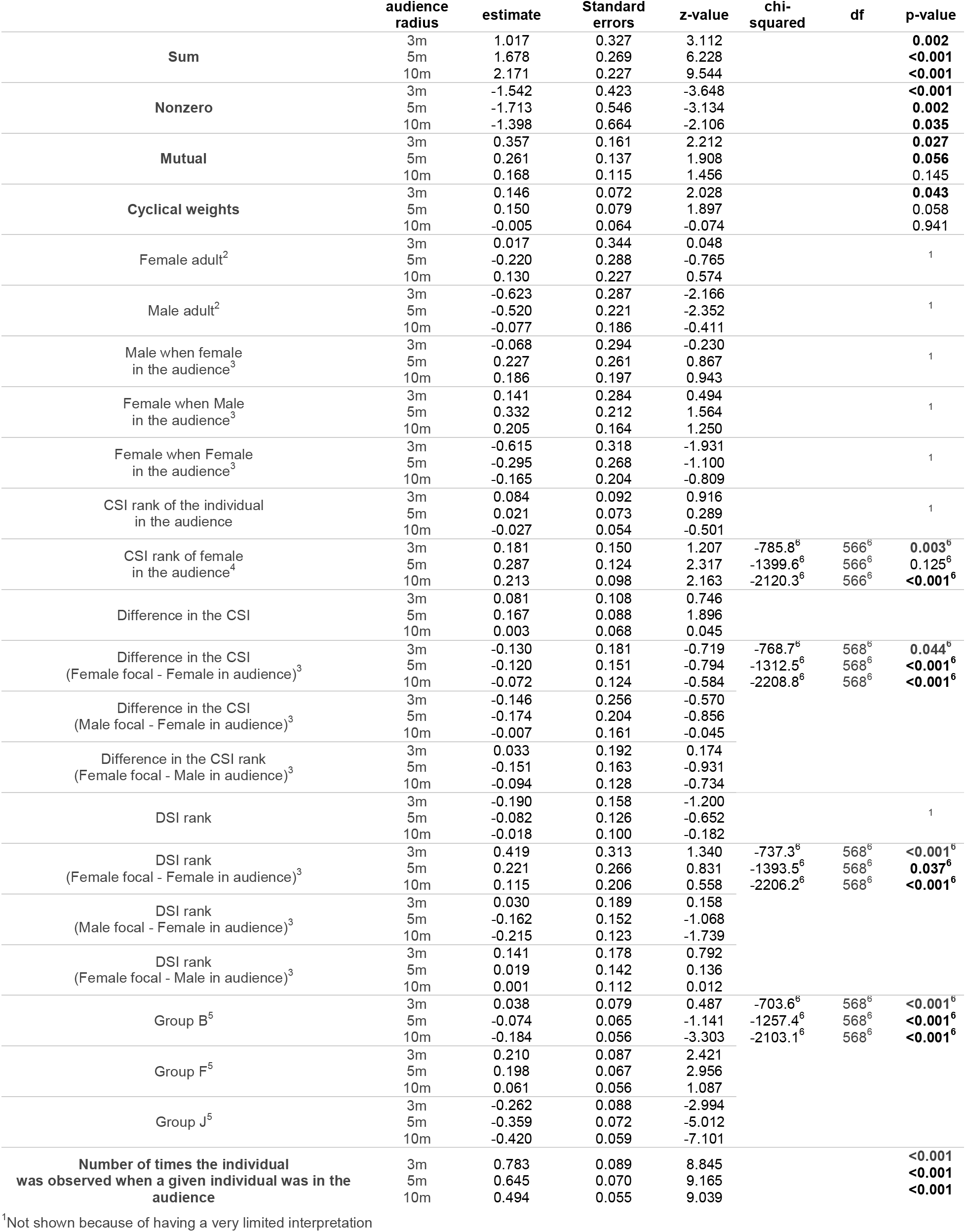
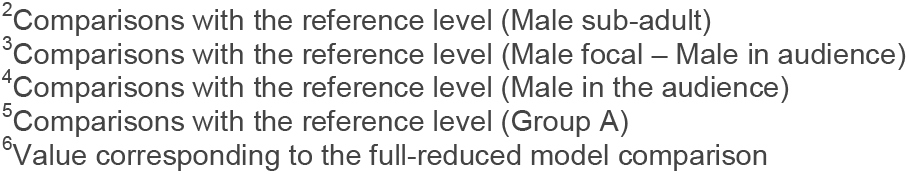
Results of the exponential graph model of the effects of audience composition within 3, 5 and 10m radius on the probability that an individual scent-mark when passing a marking spot.

|  | audience radius | estimate | Standard errors | z-value | chi-squared | df | p-value |
| --- | --- | --- | --- | --- | --- | --- | --- |
| Sum | 3m | 1.017 | 0.327 | 3.112 |  |  | **0.002** |
|  | 5m | 1.678 | 0.269 | 6.228 |  |  | **<0.001** |
|  | 10m | 2.171 | 0.227 | 9.544 |  |  | **<0.001** |
| Nonzero | 3m | -1.542 | 0.423 | -3.648 |  |  | **<0.001** |
|  | 5m | -1.713 | 0.546 | -3.134 |  |  | **0.002** |
|  | 10m | -1.398 | 0.664 | -2.106 |  |  | **0.035** |
| Mutual | 3m | 0.357 | 0.161 | 2.212 |  |  | **0.027** |
|  | 5m | 0.261 | 0.137 | 1.908 |  |  | **0.056** |
|  | 10m | 0.168 | 0.115 | 1.456 |  |  | 0.145 |
| Cyclical weights | 3m | 0.146 | 0.072 | 2.028 |  |  | **0.043** |
|  | 5m | 0.150 | 0.079 | 1.897 |  |  | 0.058 |
|  | 10m | -0.005 | 0.064 | -0.074 |  |  | 0.941 |
| Female adult<sup>2</sup> | 3m | 0.017 | 0.344 | 0.048 |  |  | 1 |
|  | 5m | -0.220 | 0.288 | -0.765 |  |  |  |
|  | 10m | 0.130 | 0.227 | 0.574 |  |  |  |
| Male adult<sup>2</sup> | 3m | -0.623 | 0.287 | -2.166 |  |  | 1 |
|  | 5m | -0.520 | 0.221 | -2.352 |  |  |  |
|  | 10m | -0.077 | 0.186 | -0.411 |  |  |  |
| Male when female in the audience<sup>3</sup> | 3m | -0.068 | 0.294 | -0.230 |  |  | 1 |
|  | 5m | 0.227 | 0.261 | 0.867 |  |  |  |
|  | 10m | 0.186 | 0.197 | 0.943 |  |  |  |
| Female when Male in the audience<sup>3</sup> | 3m | 0.141 | 0.284 | 0.494 |  |  | 1 |
|  | 5m | 0.332 | 0.212 | 1.564 |  |  |  |
|  | 10m | 0.205 | 0.164 | 1.250 |  |  |  |
| Female when Female in the audience<sup>3</sup> | 3m | -0.615 | 0.318 | -1.931 |  |  | 1 |
|  | 5m | -0.295 | 0.268 | -1.100 |  |  |  |
|  | 10m | -0.165 | 0.204 | -0.809 |  |  |  |
| CSI rank of the individual in the audience | 3m | 0.084 | 0.092 | 0.916 |  |  | 1 |
|  | 5m | 0.021 | 0.073 | 0.289 |  |  |  |
|  | 10m | -0.027 | 0.054 | -0.501 |  |  |  |
| CSI rank of female in the audience<sup>4</sup> | 3m | 0.181 | 0.150 | 1.207 | -785.8<sup>6</sup> | 566<sup>6</sup> | **0.003<sup>6</sup>** |
|  | 5m | 0.287 | 0.124 | 2.317 | -1399.6<sup>6</sup> | 566<sup>6</sup> | **0.125<sup>6</sup>** |
|  | 10m | 0.213 | 0.098 | 2.163 | -2120.3<sup>6</sup> | 566<sup>6</sup> | **<0.001<sup>6</sup>** |
| Difference in the CSI | 3m | 0.081 | 0.108 | 0.746 |  |  |  |
|  | 5m | 0.167 | 0.088 | 1.896 |  |  |  |
|  | 10m | 0.003 | 0.068 | 0.045 |  |  |  |
| Difference in the CSI (Female focal - Female in audience)<sup>3</sup> | 3m | -0.130 | 0.181 | -0.719 | -768.7<sup>6</sup> | 568<sup>6</sup> | **0.044<sup>6</sup>** |
|  | 5m | -0.120 | 0.151 | -0.794 | -1312.5<sup>6</sup> | 568<sup>6</sup> | **<0.001<sup>6</sup>** |
|  | 10m | -0.072 | 0.124 | -0.584 | -2208.8<sup>6</sup> | 568<sup>6</sup> | **<0.001<sup>6</sup>** |
| Difference in the CSI (Male focal - Female in audience)<sup>3</sup> | 3m | -0.146 | 0.256 | -0.570 |  |  |  |
|  | 5m | -0.174 | 0.204 | -0.856 |  |  |  |
|  | 10m | -0.007 | 0.161 | -0.045 |  |  |  |
| Difference in the CSI rank (Female focal - Male in audience)<sup>3</sup> | 3m | 0.033 | 0.192 | 0.174 |  |  |  |
|  | 5m | -0.151 | 0.163 | -0.931 |  |  |  |
|  | 10m | -0.094 | 0.128 | -0.734 |  |  |  |
| DSI rank | 3m | -0.190 | 0.158 | -1.200 |  |  | 1 |
|  | 5m | -0.082 | 0.126 | -0.652 |  |  |  |
|  | 10m | -0.018 | 0.100 | -0.182 |  |  |  |
| DSI rank (Female focal - Female in audience)<sup>3</sup> | 3m | 0.419 | 0.313 | 1.340 | -737.3<sup>6</sup> | 568<sup>6</sup> | **<0.001<sup>6</sup>** |
|  | 5m | 0.221 | 0.266 | 0.831 | -1393.5<sup>6</sup> | 568<sup>6</sup> | **0.037<sup>6</sup>** |
|  | 10m | 0.115 | 0.206 | 0.558 | -2206.2<sup>6</sup> | 568<sup>6</sup> | **<0.001<sup>6</sup>** |
| DSI rank (Male focal - Female in audience)<sup>3</sup> | 3m | 0.030 | 0.189 | 0.158 |  |  |  |
|  | 5m | -0.162 | 0.152 | -1.068 |  |  |  |
|  | 10m | -0.215 | 0.123 | -1.739 |  |  |  |
| DSI rank (Female focal - Male in audience)<sup>3</sup> | 3m | 0.141 | 0.178 | 0.792 |  |  |  |
|  | 5m | 0.019 | 0.142 | 0.136 |  |  |  |
|  | 10m | 0.001 | 0.112 | 0.012 |  |  |  |
| Group B<sup>5</sup> | 3m | 0.038 | 0.079 | 0.487 | -703.6<sup>6</sup> | 568<sup>6</sup> | **<0.001<sup>6</sup>** |
|  | 5m | -0.074 | 0.065 | -1.141 | -1257.4<sup>6</sup> | 568<sup>6</sup> | **<0.001<sup>6</sup>** |
|  | 10m | -0.184 | 0.056 | -3.303 | -2103.1<sup>6</sup> | 568<sup>6</sup> | **<0.001<sup>6</sup>** |
| Group F<sup>5</sup> | 3m | 0.210 | 0.087 | 2.421 |  |  |  |
|  | 5m | 0.198 | 0.067 | 2.956 |  |  |  |
|  | 10m | 0.061 | 0.056 | 1.087 |  |  |  |
| Group J<sup>5</sup> | 3m | -0.262 | 0.088 | -2.994 |  |  |  |
|  | 5m | -0.359 | 0.072 | -5.012 |  |  |  |
|  | 10m | -0.420 | 0.059 | -7.101 |  |  |  |
| Number of times the individual was observed when a given individual was in the audience | 3m | 0.783 | 0.089 | 8.845 |  |  | **<0.001** |
|  | 5m | 0.645 | 0.070 | 9.165 |  |  | **<0.001** |
|  | 10m | 0.494 | 0.055 | 9.039 |  |  | **<0.001** |

<sup>1</sup>Not shown because of having a very limited interpretation
- <sup>2</sup>Comparisons with the reference level (Male sub-adult) <sup>3</sup>Comparisons with the reference level (Male focal – Male in audience) <sup>4</sup>Comparisons with the reference level (Male in the audience) <sup>5</sup>Comparisons with the reference level (Group A) <sup>6</sup>Value corresponding to the full-reduced model comparison

**Figure 2:**
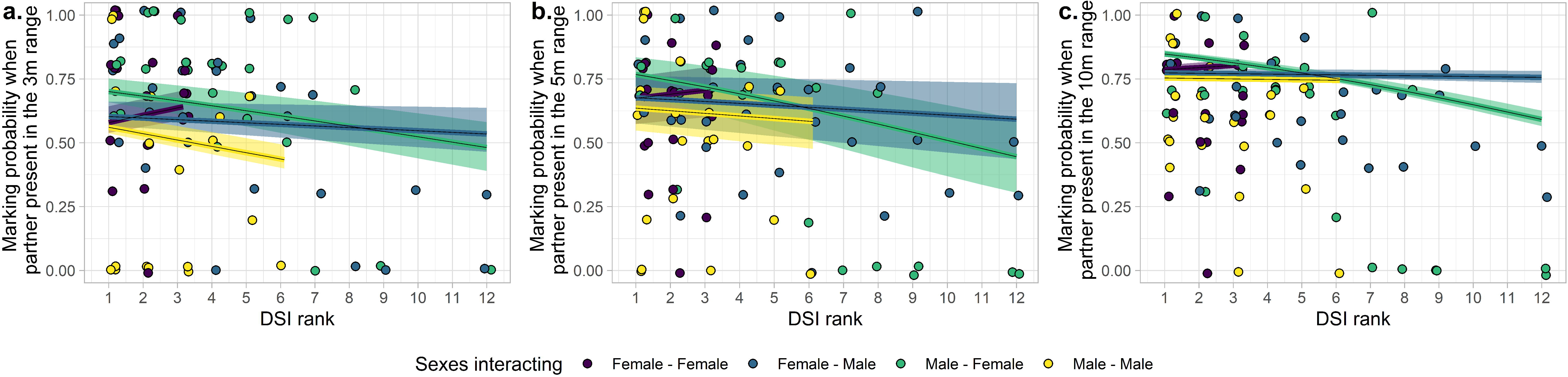
Probability to scent-mark as a function of the DSI rank of the dyad for an audience a) in a 3m radius, b) in a 5m radius, and c) in a 10m radius. The shaded areas show 95% confidence intervals of the model (conditional on the number of observations being at its average and on a group effect weighted by the number of individuals in each group).

There was a significant interaction effect between the sexes and the CSI difference between individuals of the respective dyad within the 3m and 10m radius (full-reduced model comparison: 3m chi2=-740.8, df=568, p-value=0.004; 10m chi2=-2225.5, df=568, p-value<0.001; Table 4) but not within the 5m radius (chi2=-1393.5, df=568, p-value=0.152). Males scent-marked more often in front of females that were more social than themselves (i.e. when the CSI difference was negative; Fig.3). Females scent-marked more often in front of females that were more social than themselves (Fig.3).

**Figure 3:**
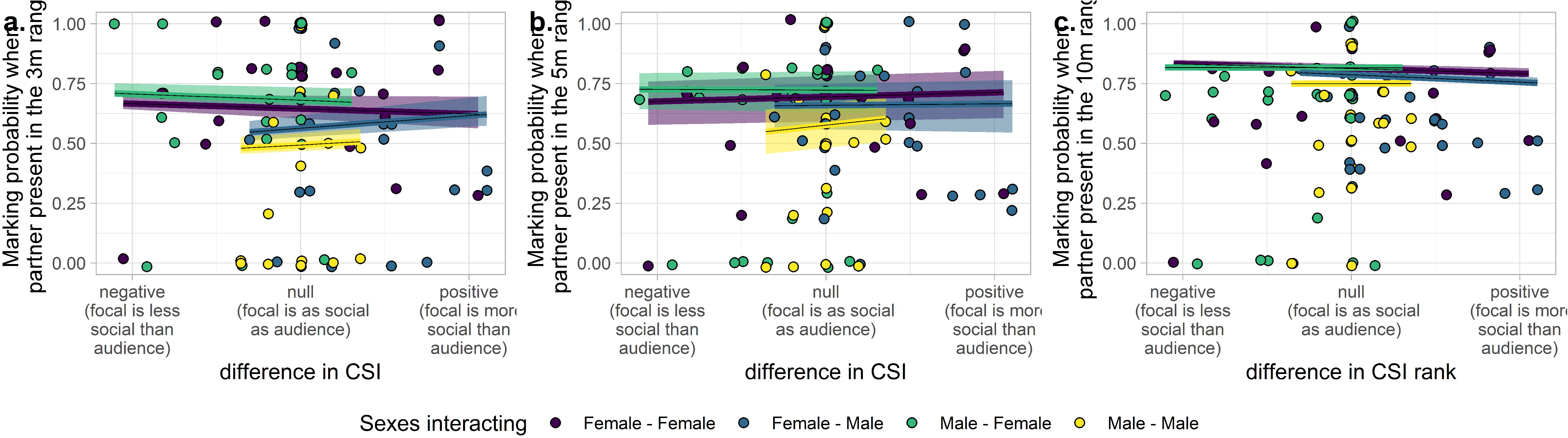
Probability to scent-mark as a function of the difference in CSI of the dyad for an audience a) in a 3m radius, b) in a 5m radius, and c) in a 10m radius. The shaded areas show 95% confidence intervals of the model (conditional on the number of observations being at its average and on a group effect weighted by the number of individuals in each group).

There was also a significant interaction effect between sex and CSI rank of the individual in the audience (full-reduced model comparison: 3m chi2=-776.0, df=566, p-value<0.001; 5m chi2=-1351.9, df=566, p-value<0.001; 10m chi2=-2237.0, df=566, p<0.001; Table 4). More specifically, individuals scent-marked more often in front of the less social females (the ones exhibiting a greater CSI rank; Fig.4).

**Figure 4:**
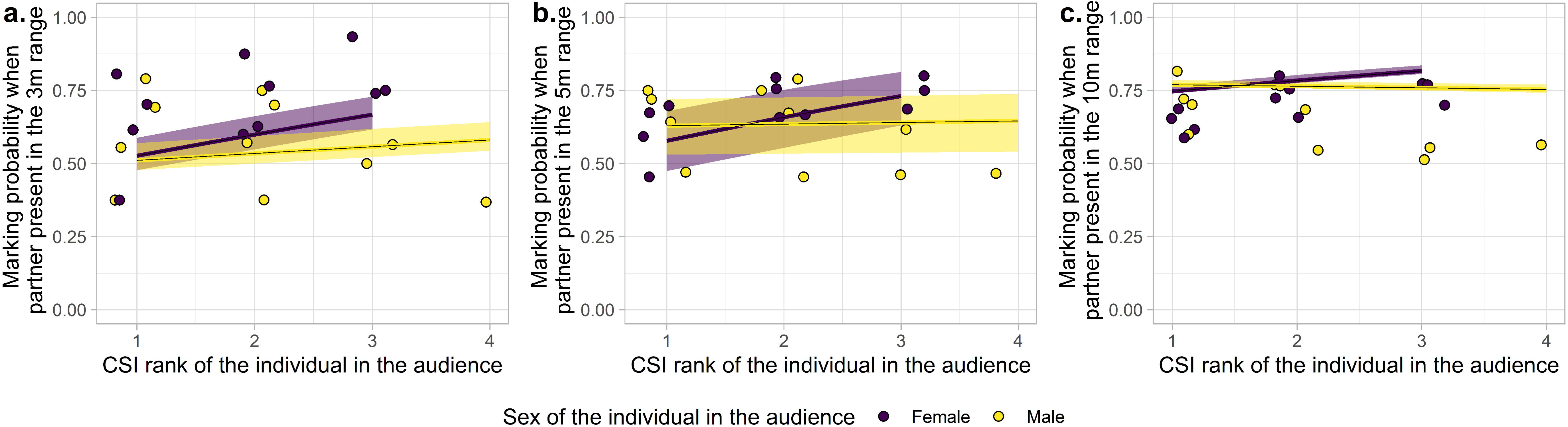
Probability to scent-mark as a function of the CSI rank of the individual in the audience when a) in a 3m radius, b) in a 5m radius, and c) in a 10m radius. The shaded areas show 95% confidence intervals of the model (conditional on the number of observations being at its average and on a group effect weighted by the number of individuals in each group).

The probability for an individual to be observed marking in front of a given conspecific increased as a function of the number of times the individual was observed in the presence of this individual (3m: b= 0.783, p<0.001; 5m: b=0.645, p<0.001; 10m: b=0.494, p<0.001; Table 4). At 3m and 5m, the probability that the audience effect would be reciprocate was significantly higher than the probability that the audience effect is not reciprocated (mutual term: b=0.357, p=0.027 and b=0.261, p=0.056 respectively; Table 4). This effect was not significant at a 10 m radius (b=0.168, p=0.145; Table 4). There was a tendency towards nonlinear hierarchical structure of the network at 3m (cyclical clustering term: b= 0.146, p=0.043; Table 4) but this effect no longer existed at 5 and 10m (b=0.150, p=0.058 and b=-0.005, p=0.941 respectively; Table 4). In addition, there was a significant effect of the group on the probability to observe scent-marking (full-null model comparison: 3m: chi2=-729.6, df=568, p<0.001; 5m: chi2=-1222.2, df=568, p<0.001; 10m: chi2=-2122.1, df=568, p<0.001; Table 3).

## Discussion

In this study, we investigated intra-group audience effects on anogenital scent-marking in a wild population of redfronted lemurs. Our results indicate that scent-marking in redfronted lemurs is associated with some behavioural flexibility linked to the composition of the audience at the time of scent deposition. Moreover, our results also show that the nature of the audience effects observed differed between males and females, with males being more sensitive to their audience than females.

On the intersexual level, we predicted that anogenital scent-marking in males might function as a form of mate attraction or advertisement with males performing more anogenital scent-marking in the presence of females and that this effect should be dependent on the social relationships the male maintains with the females present in the audience. Males were observed to scent-mark more in front of a female when they had a stronger relationship with that female (lower DSI rank). Males also seem to scent-mark more often in front of the most social females of their group. These two results together suggest that males may address scent-mark signals to females with whom they maintain a close relationship but that are also engaged in close social relationships with other individuals of their group. This result is coherent with the literature showing that the most social males are the ones scent-marking the most^58^. In contrast, our results revealed no effect of the male audience on the probability of females to anogenital-mark. Hence, as predicted, in redfronted lemurs, anogenital scent-marking may have a mate attraction or advertisement function only in males, as also shown in lemur species exhibiting female dominance (e.g. *P. verreauxi*^65^, *L. catta*^28^, Grey mouse lemur, *Microcebus murinus*^66^). Hence, on the intersexual level, the dominance structure does not appear to predict the function of anogenital scent-marks in males across lemurs.

On the intrasexual level, we predicted that both males and females should modify their scent-marking behaviour in response to a same-sex audience and that this effect should be particularly dependent on their social value and the one of the individuals in the audience. Males were observed to scent-mark significantly less often when a greater proportion of males of their group were within the 3m radius. This observation is reinforced by the lowest anogenital scent-marking probabilities being associated with the male-male category in the outputs of the exponential random graph analyses (Fig.2 and 3). The probability that a male would scent-mark in front of another male decreased when these two males had a weaker social relationship (greater DSI rank). Males also tended to mark less often in front of males that were more social than themselves (positive values of the difference in CSI). Hence, males seem to avoid scent-marking in close proximity with an increased number of males and in the presence of a male with whom they have a weak affiliation, especially if the latter is more social than themselves. Even if there is no linear hierarchy and low aggression levels in male redfronted lemurs^58^, the risk of physical aggression might be elevated when scent-marking in close proximity. In contrast, males that are further away might either be less motivated or less successful in suppressing another male scent-marking in the five- or ten-meters radius. Although we do not know whether males can suppress scent-marking in other males, investigations on the probability of exhibiting aggression at different distances might help to elucidate potential mechanisms. If males are more likely to engage in aggression when they are in 3m in comparison to 5m or 10m, avoidance of potential aggression might explain the suppression of scent-marking behaviour in males. Alternatively, some males may give priority to other males to scent-mark the spot when they are in proximity. Hence, competition among males may result in having priority of access to these specific scent-marking spots. If the male to whom the priority is given is at a distance of 5 or 10m, the focal male might still have time to scent-mark before its arrival. Our results indicate that priority of scent-marking seems to be given to the most social males, which may also contribute to explain why the central males have been observed to be the ones scent-marking more frequently^58^. Hence, in redfronted lemurs, males might use anogenital scent-marking as a way to advertise their social status to other males as an indirect form of competition as it has also been suggested for other lemur species (*Lemur catta*^7,69^; *Propithecus verreauxi*^50,68,72,76^; *Eulemur rufus*^33^; *P. edwardsi*^67,77^*; P. candidus*^67^).

The highest scent-marking probabilities are associated with the female-female category in the outputs of the exponential random graph analyses (Fig.2, 3), suggesting that overall, females seem to be less sensitive to the presence of females in their audience than to the presence of males and that males seem to be overall more sensitive to their audience than females. In females, we found an audience effect, with the probability that a female scent-marked in front of another female increasing when these two females had a weaker social relationship (greater DSI rank). Moreover, we found a trend for individuals to scent-marked more often in front of the less social females (the ones exhibiting a greater CSI rank). Hence, despite the absence of hierarchical dominance instigated through overtly aggressive behaviour, females may signal their social status to other females; in particular, the most social ones may address these signals to the less social ones.

Overall, our results indicate that males seem to be more sensitive than females to their audience when scent-marking. On the intersexual level, only males seem to be sensible to the audience composition. On the intrasexual level, scent-marks seem to have a social status signalling function in both sexes as it has been suggested for other congeners from groups lacking clear dominance patterns both between and within sexes^51^. Studying the flexibility of complex signal usage across social contexts (audience compositions) contributes to uncovering the particular social characteristics eliciting or constraining complex signal expression ^78,79^. These social characteristics may, in turn, constitute social pressures acting for or against the evolution of complex signalling behaviours^24,78,80,81^. Hence, in species without overt dominance relations, males seem to be more constrained in the expression of scent signals and to adjust their scent-marking behaviour in a more fine-tuned manner to the composition of the audience than females. Less social males, which scent-mark less frequently in the presence of other males, may have to rely primarily on the long-lasting component of the signal to advertise their social status, thereby avoiding potential aggression from other males. Overall, our results are concordant with the observations that genital and perianal secretions are chemically more complex in egalitarian species living in multi-male-multi-female groups than in species living in pairs and/or having female dominance and that males have more chemically complex genital and perianal secretions than females in these species without overt dominance relations^36,48^.

Studying audience effects is an interesting gateway to distinguish between arousal-mediated and intentional communication^81,82^. Audience effects may indeed reveal a potential intentional communication, primarily when this variation is based on subtle social and behavioural variations such as the quality of relationships^81–85^. Here we show that redfronted lemurs do not only flexibly scent-mark as a function of the number of individuals present in the audience but also based on the strength of social relationships they maintain with specific individuals present in the audience. Such social competence (referred to as ‘social use’) was described as one indicator of potential voluntary control of signalling behaviour^82,86^.

Finally, some limitations and possible factors increasing the noise in our observations need to be mentioned. First, some individuals may also choose not even to pass a specific scent-marking spot in the presence of a particular audience. Hence, we cannot exclude and control for a potential audience effect on the probability to pass this spot or not. Second, the effect of who may have marked beforehand on a specific spot may also be highly relevant in an individual’s choice to mark or not when passing a spot. This parameter is hard to control in the field because we do not have information on the possible passage on this spot before the observation and video recording started (and olfactory signals are long-lasting). Further studies on the patterns of scent-marking behaviour succession occurring on a given scent-marking spot may uncover these questions. Moreover, considering the directionality of the individuals in the audience may also be an interesting perspective in this regard. While at three meters, the individuals may be relatively homogeneously attentive to the scent-marking of an individual, at five and ten meters, the directionality of the individuals in the audience may be more important to consider. Individuals approaching the scent-marking spot may indeed be more attentive than individuals that already overpassed this spot. This may also contribute to the loss of some audience effects observed at larger distances.

Besides these intra-group functions, scent-marking may also be a form of inter-group communication, with a function of resource or territorial defence through individual or group odour deposition^37,87–89^. Female red-fronted lemurs are philopatric and inherit the territory of their mother, so they may have an especially steep interest in defending their territory and/or its associated resources. As some of the events reported here occurred in a context of post or pre-inter group encounter (with no extra-group individuals in the audience), this could have an impact in our results, however, the effect of context was not significant when looking at the effect of the proportion of males and females present in the audience. Hence, exploring in more detail inter-group audience effects may reveal interesting information to complete the full picture of anogenital scent-marking functions in redfronted lemurs.

In conclusion, we showed that scent-marking in redfronted lemurs is associated with some behavioural flexibility linked to the composition of the audience (i.e. number and social value of the individuals present), ascribing redfronted lemurs social competence in this context. Moreover, our approach broadens our understanding of signal delivery and its associated sex differences, providing important insights into the functional significance of anogenital scent-marking in redfronted lemurs. As such, this study offers a potential behavioural pattern associated with anogenital scent-marking in species without overt dominance relationships that exhibit similarities but also differences to those described for species exhibiting female dominance, supporting the notion that the social systems co-varies with scent-marking behaviours and scent-complexity in strepsirrhines.

## Material and methods

### Study Site and Subjects

We conducted this study in Kirindy Forest, a dry deciduous forest located ca. 60 km north of Morondava, western Madagascar, managed within a forestry concession operated by the Centre National de Formation, d’Etudes et de Recherche en Environnement et Foresterie (CNFEREF)^47^. Since 1996, all members of a local population of redfronted lemurs inhabiting a 70-ha study area within the forest have been regularly captured, marked with individual nylon or radio collars, and subjected to regular censuses and behavioural observations as part of a long-term study^47^. The data presented in this study were collected from May to November 2018 on 28 adult individuals belonging to four groups (11 females and 17 males). Among males, 14 were adults and 3 sub-adults (1.5-2 years). Sub-adults were included in the study as they were observed to perform scent-marking behaviours as often as adult individuals. Reproduction of the species is seasonal, with a 4-week mating season in May–June and a birth season in September–October^58,90^. All applicable international, national, and/or institutional guidelines for the care and use of animals were followed. The authors complied with the ARRIVE guidelines. This study adhered to the Guidelines for the Treatment of Animals in Behavioral Research and Teaching^91^ and the legal requirements of the country (Madagascar) in which the work was carried out. The protocol for this research was approved by the Commission Tripartite de la Direction des Eaux et Forêts (Permit No 47 and 215 18/MEF/SG/DGF/DSAP/SCB.Re).

### Data Collection

Between May to July (later referred to by mating season) and September to November (later referred to as birth season), data were collected by focal scent-mark observations. Scent-marking behaviours were observed *ad libitum* during 27 to 34 half-days in each group. During these sessions, a total of 120 scent-marking behaviours (26 to 34 per group) served as foci for 15 minutes observations that were video recorded. During these 15 minutes observation periods, we annotated each individual passing the focal scent-mark spot, its identity, whether it performed scent-marking or not, the date, the time, the context and the identity of all the other individuals present in the radius of 3, 5 and 10 meters. The context was classified using four categories: resting, feeding, travelling and disturbance defining the group activity. The context ‘disturbance’ referred to situations in which individuals of the group are vigilant, and none of the other three context categories could be attributed to the situation. Cases when individuals of another group were visible were excluded.

Additionally, from May to November 2018, we also carried out 30 minutes individual focal observations in the morning between ca. 07:00–10:00 h and afternoon between14:00–17:00 h. A given individual was never observed for more than one 30 minutes session per day, and observations were balanced among each observation hour for each individual. The final dataset included 367 hours of focal observations, with an average of 14.7 hours per individual.

### Data analyses

All analyses were carried out using R (version 3.6.0)^92^ and RStudio (version 1.2-1335)^93^.

#### Social values of individuals

We calculated the CSI (Composite Sociality Index^94^; equation (1)) for each individual based on three mutually exclusive affiliative behaviours: body contact, grooming and huddling. For each individual, we first calculated the proportion of time spent in body contact, huddling and grooming with an individual of its group (except juveniles). The resulting hourly rates for each of the three behaviours (***r. bc***_***i***_, ***r***. ***hu***_***i***_, ***r***. ***gr***_***i***_) were next divided by the respective mean rate for the group of the given individual before being summed up. To obtain the CSI, the summed value was divided by three, corresponding to the number of behaviours considered. We further attributed to each individual a CSI rank within each group and age-sex category, with individuals of rank 1 being the ones interacting the most often. To obtain the difference in CSI between two individuals we subtracted the CSI value of the individual in the audience to the one of the focal. These CSI difference values were scaled using the R function ‘scale’ within each group.

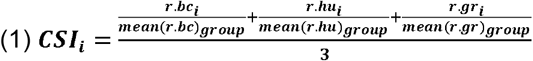

#### Social values of dyads

We calculated the DSI (Dyadic Composite Sociality Index^94^; equation (2)) of each dyad of individuals in a given group (excluding juveniles) following the same principle as for the CSI. Because two individuals were never observed simultaneously in a given group, interaction rates for a given dyad A-B could be calculated by summing up the rates associated with A being focal and interacting with B and B being focal and interacting with A. For each individual, we first calculated the time spent in body contact, huddling and grooming with each of its adult group members and divided it by the total observation duration of this individual while its partner was present in the group. We further attributed to each dyad a DSI rank within each group and age-sex category, with dyads of rank 1 being the most social dyads of their group. We used rank instead of raw DSI as we were not interested in group differences. In this way, the most social dyad of each group is attributed with the same social value.

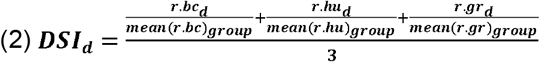

#### Estimation of the audience effect on anogenital scent-marking

For a given individual, we only considered anogenital marking events that occurred with a time-lapse of at least 5 minutes between each other. We selected passing events (without scent-marking) on the same criteria. We included only individuals for whom we had at least two observations of each passing and marking. Three males that emigrated during the period had to be excluded because we had only one observation of either passing or marking. The final male dataset included 14 individuals (3 sub-adults and 11 adults) observed for 60 pass events and 105 anogenital marking events. The female dataset included 11 adult females observed for 44 pass-by and 118 scent-marking events.

We first ran two independent Generalized Linear Mixed Models (GLMM) for both sexes, estimating the influence of the audience composition on the probability of anogenital-marking behaviour to occur at a given time. These models had a binomial error structure and logit link function^95^ and were run for each audience radius. These models were fitted using the function glmer of the R package lme4 (version 1.1-21)^96^ with the optimiser’ ‘bobyqa’. As fixed effect, we included the proportions of males and adult females present in the given distance radius. To control for age (for males only as we only had one age class for females), context and season we also included these terms in the model as control predictors. Individual identity and date were included as random factors to account for individual variations and the possible effect of particular events.

To reduce the risk of type I errors^97^, we included all possible random slopes components (the proportion of males, the proportion of adult females, context and season within individual identity). We manually dummy-coded and centred context, season and age, and z-transformed the proportion of males and the proportion of females before including them as random slopes. Initially, we also included all correlations among random intercepts and slopes for all models. However, for females, these were all estimated to have absolute values being essentially one indicating that they were not identifiable^98^. Hence, we removed these correlations from the female model.

As an overall test of the effect of audience composition on the probability to anogenital scent-mark, we compared the full model with the null model lacking the fixed effects characterising the audience (proportion of males and proportion of females) but comprising the control fixed effects and the same random effect structure as the full model^97^. This comparison was performed using a likelihood ratio test^99^.

Model stability was assessed by comparing the estimates of the model run on the full dataset with the ones run on datasets, excluding each level of the random effects one after the other^100^. The models were relatively stable (for males: Supplementary File S1.A; for females: Supplementary File S1.C). To control for potential collinearity problems, we calculated the Variance Inflation Factors^101^ for the model, excluding the random effects. VIF values ranged from 1.03 to 1.76 for the males (Supplementary File S1.B) and from 1.07 to 2.20 for females (Supplementary File S1.D).

Confidence intervals were derived using the function bootMer of the package lme4, using 1,000 parametric bootstraps and bootstrapping over the random effects, too (argument’ use.u’ set to TRUE). Tests of the individual fixed effects were derived using likelihood ratio tests^102^ (R function drop1 with argument’ test’ set to” Chisq”). We determined the proportion of the total variance explained by the fixed effects (R^2^_m_; marginal coefficient of determination), and the proportion of the variance explained by both fixed and random effects (R^2^_c_; conditional coefficient of determination) following the method recommended by Nakagawa et al.^103^ and using the function r.squaredGLMM of the package MuMIn (version 1.43.6)^104^. Because our models seem to suffer singularity issues, we further applied a Bayesian method as recommended by the authors of the "lme4" package^96^. This approach should allow both regularising the model via informative priors and giving estimates and credible intervals for all parameters that average over the uncertainty in the random-effects parameters. Details on the methods and outputs of these models are provided in supplementary material (Supplementary File S2).

To account for the nonindependence of individuals within a group and the network structure of their interactions, we first used valued exponential random graph models (ERGM)^105^ to understand how the nature of the audience may influence the probability of anogenital marking. We implemented an ERGM based on a directional weighted matrix corresponding to the number of observed anogenital marking events of a focal individual (tail) when a given individual of its group was in the audience (head).

Models were implemented with a Poisson reference distribution, and the term “sum” corresponding to the sum of the edge weights (equivalent to an intercept in a linear modelling scenario) was added to the model. In addition, a “nonzero” term was added to control for zero inflation in the distribution of edge weights. Moreover, because structural terms are essential for correct model specification^106,107^ we included a mutuality term (sum of the minimum edge weights for each potential edge), and a cyclical weights term allowing for exploring hierarchical structure^108^. Two terms were included as control predictors: an edge covariate term to account for the amount of time an individual was observed in the presence of a given individual in the audience^109^ and a node-level covariate term to control for the effect of the group. Moreover, an offset term was added to acknowledge the fact that we only consider intra-group interactions. The terms described so far were the terms remaining in the null-model.

As node level predictors, we included the interaction between sex (only for adults) and the CSI rank of the individual in the audience (in-edges). As edge covariates, we included the interaction between the sexes and the difference between the CSI of the focal individual and the individual in the audience and the interaction between the sexes and the DSI rank corresponding to the dyad in question. All the terms corresponding to the main effects and the dummy variables (with the exception of the reference male-male) were also included in the model.

ERGMs were implemented in R using the statnet suite of packages^105,110–113^. The code to implement this model is provided in the ESM, Supplementary File S3. We manually dummy coded and centred the sex interacting and z-transformed all the explanatory variables before including them into the model. The goodness of fit was assessed for each model by simulating 1000 networks and comparing the distribution of their coefficients to the observed coefficients^114,115^(Supplementary Fig. S1, S2 and S3). MCMC diagnostics were used to assess ERGM convergence (“mcmc.diagnostics” function in the ergm package) (Supplementary Fig. S4, S5 and S6). To assess the overall test of the significance of the interaction between sex and sociality we compared the deviance of the full model to the one of the null model described above. This comparison was based on a likelihood ratio-test^97,99^, R function anova with the argument test set to “chisq”. To test the significance of the individual interactions between sex and the three social variables, we compared the full model’s deviance with that of a corresponding reduced model not comprising this interaction. Confidence intervals for the interaction effects were obtained by bootstrapping the response matrix (adding or subtracting 1 to an intra-group edge weight).

## Acknowledgements

We warmly acknowledge Dr Pavel N. Krivitsky for his reactivity and help with the ERGM implementation. We are also thankful to Dr Franziska Hübner for her relevant comments on the data analyses. We are most grateful to the local team of the Kirindy field station. We thank the Malagasy Ministère de l’Environnement et des Eaux et Forêts, the Département de Biologie Animale of Antananarivo University, and the Centre National de Formation, d’Etudes et de Recherche en Environnement et Foresterie for supporting and authorising our long-term research in Kirindy. This study was funded by grants by the Deutsche Forschungsgemeinschaft (DFG FI 929/12-1 and KA 1082/35-1).

## Author contributions

L.R.P., C.F. and P.M.K conceptualised this project, L.R.P. and L.S.M developed the methodology, L.R.P. and A.M. collected data in the field, L.R.P. analysed the data, L.R.P. drafted the MS, and all authors participated in reviewing and editing the MS.

## Additional information

### Data availability

The datasets generated and analysed during the current study are available from the corresponding author on reasonable request.

### Ethics declarations

All applicable international, national, and/or institutional guidelines for the care and use of animals were followed. The authors complied with the ARRIVE guidelines. This study adhered to the Guidelines for the Treatment of Animals in Behavioral Research and Teaching^91^ and the legal requirements of the country (Madagascar) in which the work was carried out. The protocol for this research was approved by the Commission Tripartite de la Direction des Eaux et Forêts (Permit No 47 and 215 18/MEF/SG/DGF/DSAP/SCB.Re).

