## Supplementary Figure S1 for "Males are more sensitive to their audience than females when scent-marking in the redfronted lemur"

**Supplementary Figure S1:** goodness of fit for the exponential random graph model on the audience effect on scent-marking in redfronted lemurs when considering a 3m radius. For each model term, the estimate of the model (blue lines) is compared to the distribution of the estimates of 1000 simulated networks.

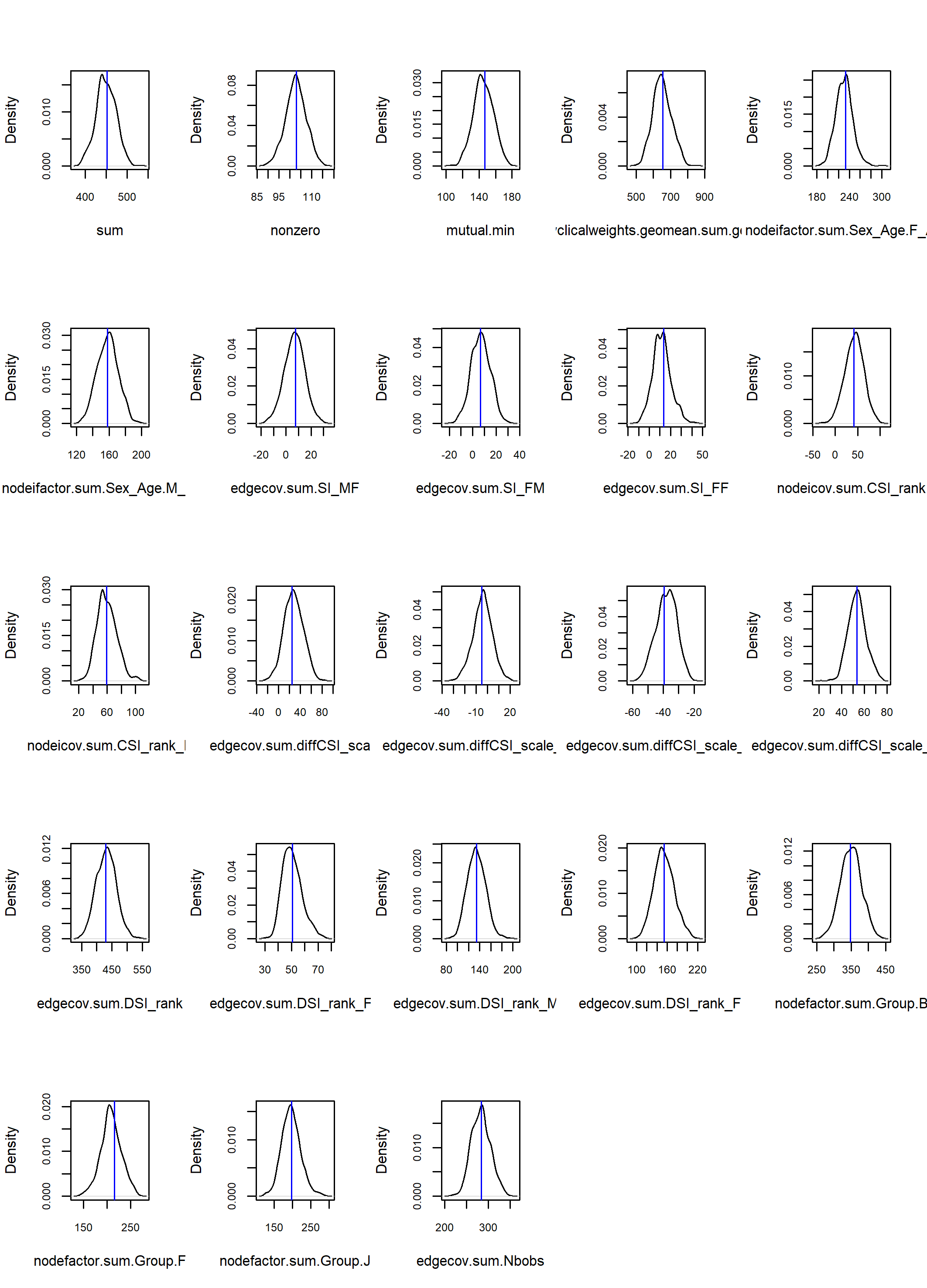
