## Supplementary Figure S4 for "Males are more sensitive to their audience than females when scent-marking in the redfronted lemur"

**Supplementary Figure S4:** MCMC diagnostics for the exponential random graph model on the audience effect on scent-marking in redfronted lemurs when considering a 3m radius. For each model term, in the first column figures illustrate Markov chains for the parameter, α; in the second column are given posterior distributions for α.

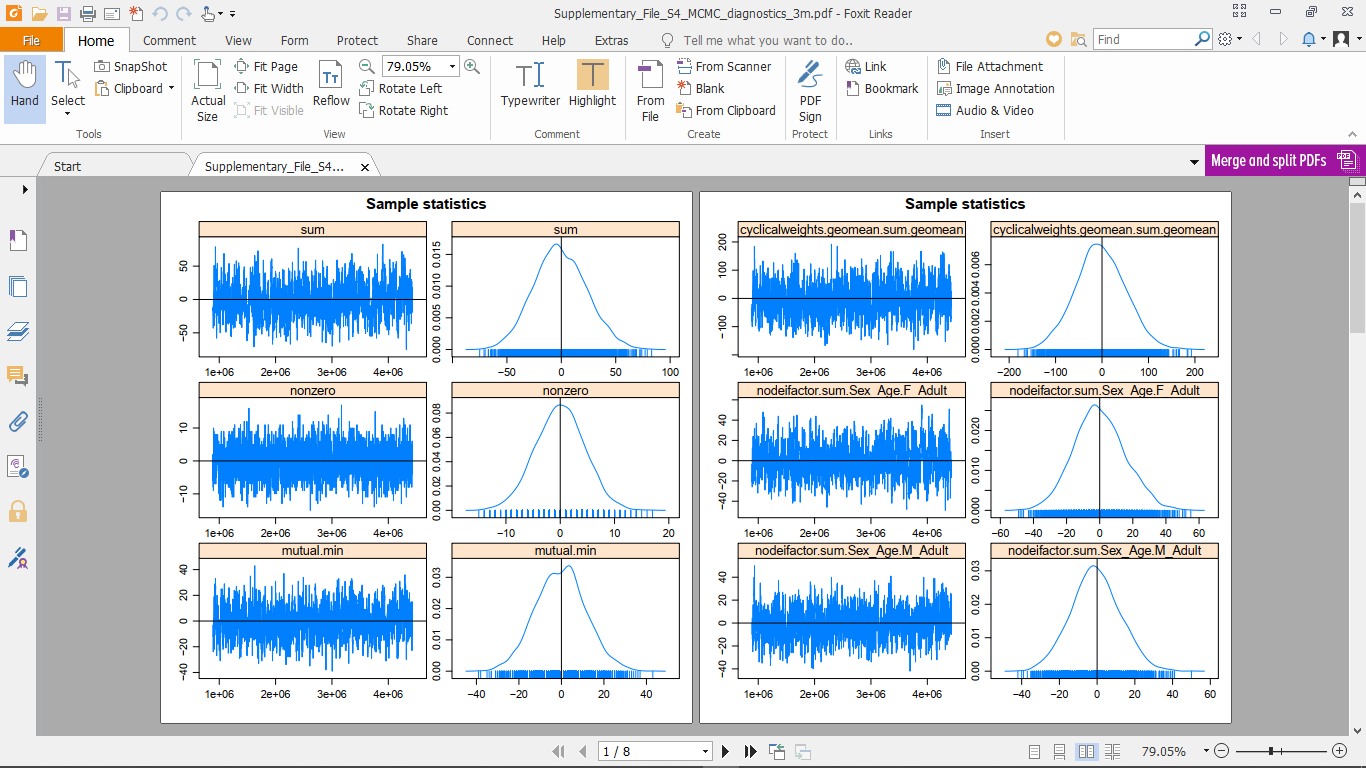

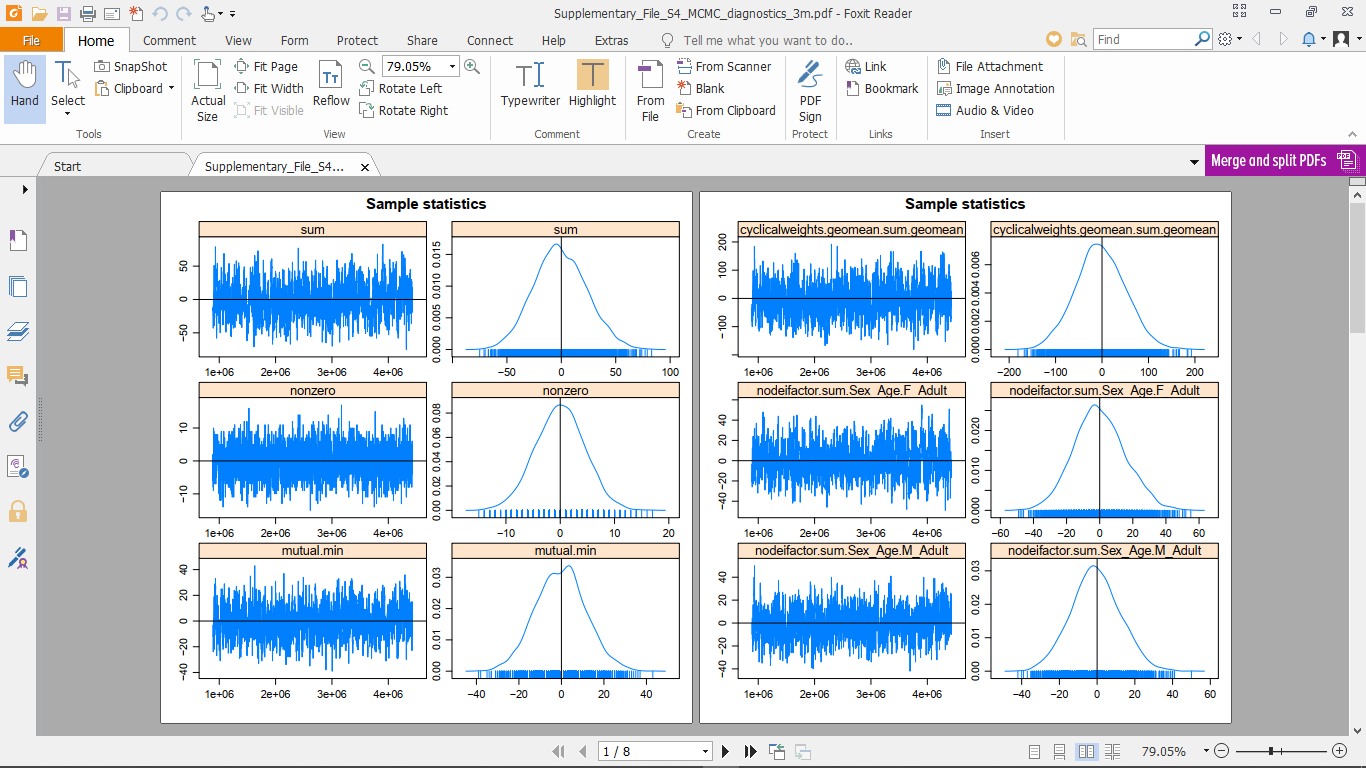

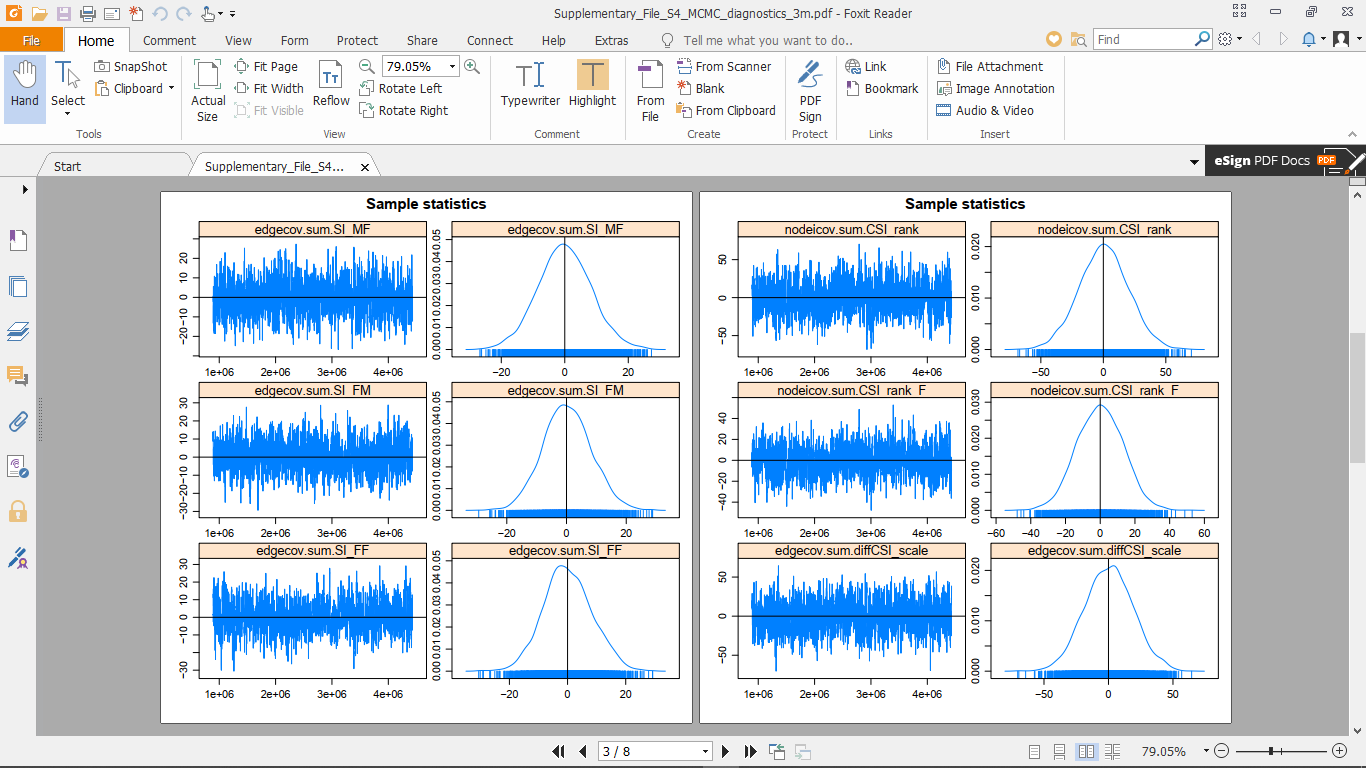

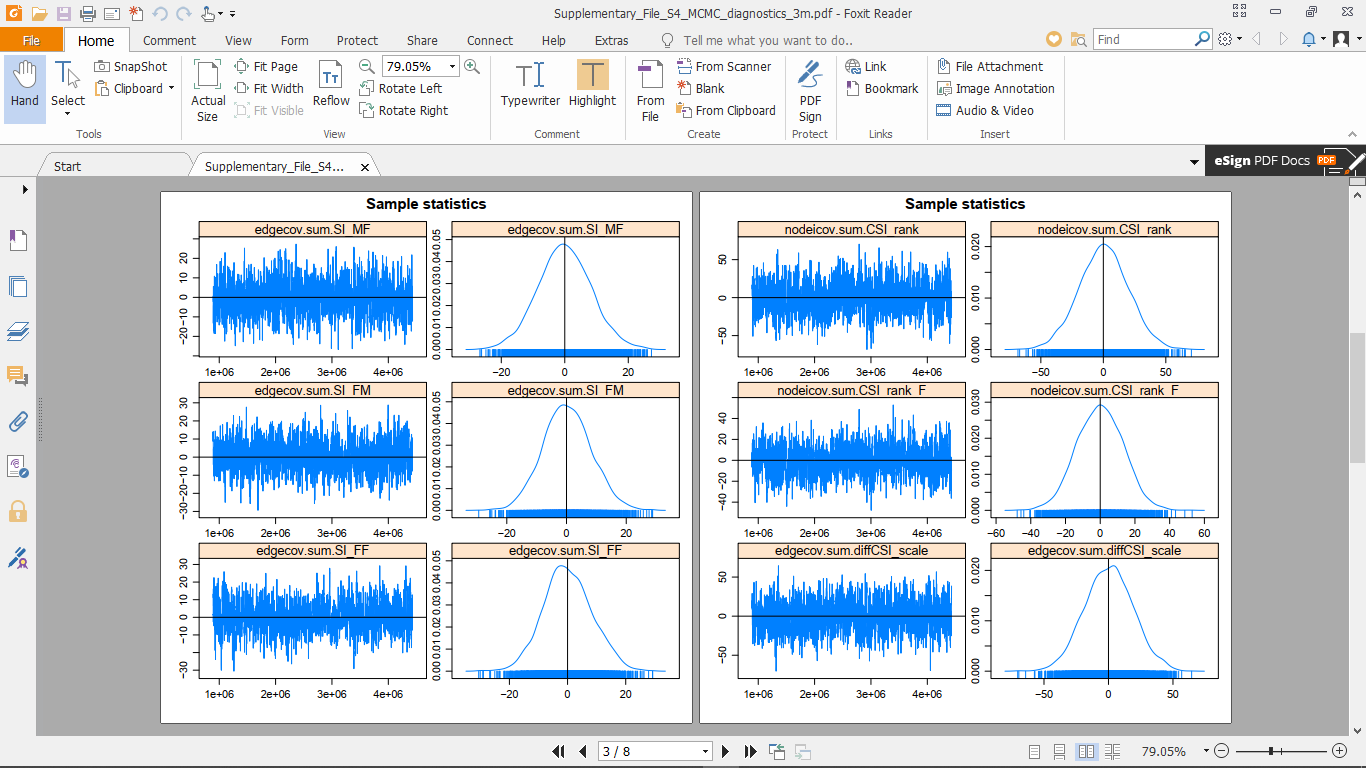

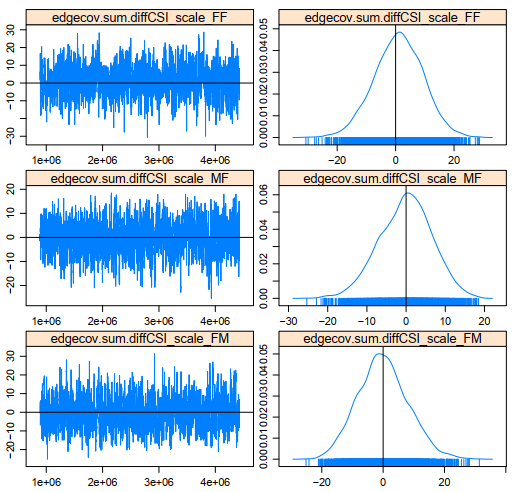

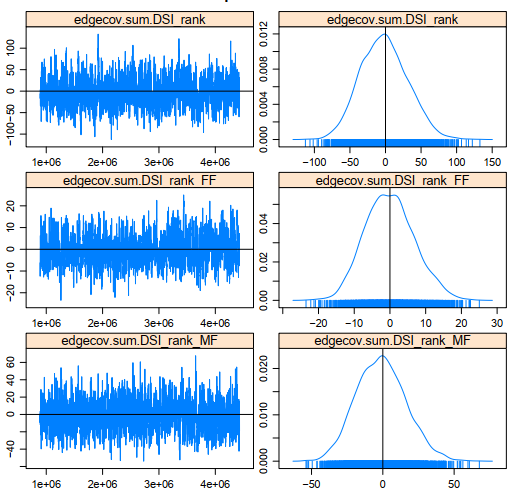

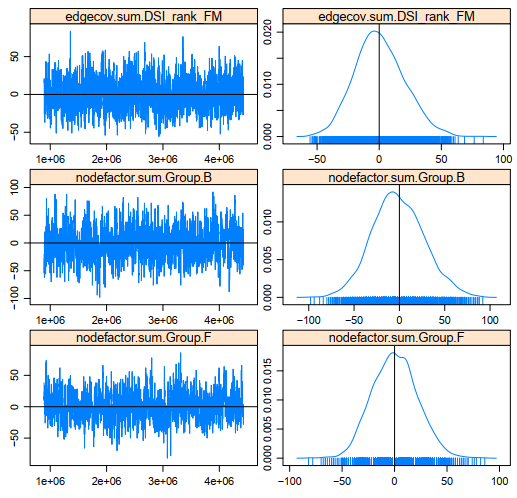

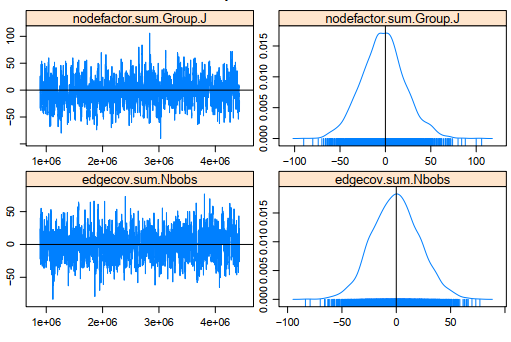
