## Supplementary Figure S5 for "Males are more sensitive to their audience than females when scent-marking in the redfronted lemur"

**Supplementary Figure S5:** MCMC diagnostics for the exponential random graph model on the audience effect on scent-marking in redfronted lemurs when considering a 5m radius. For each model term, in the first column figures illustrate Markov chains for the parameter, α; in the second column are given posterior distributions for α.

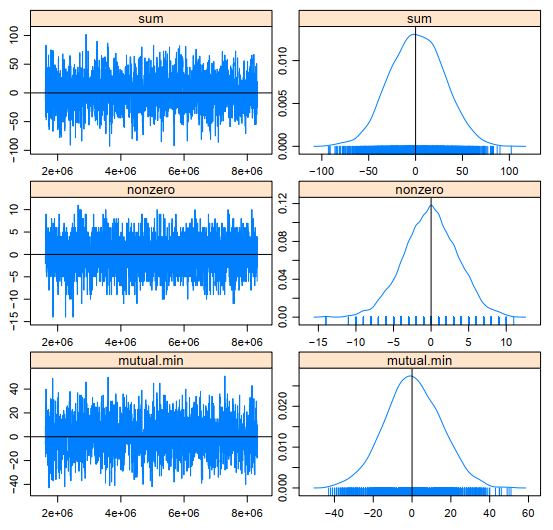

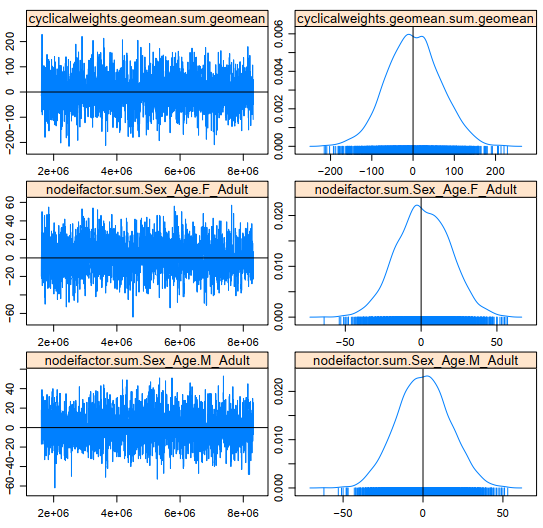

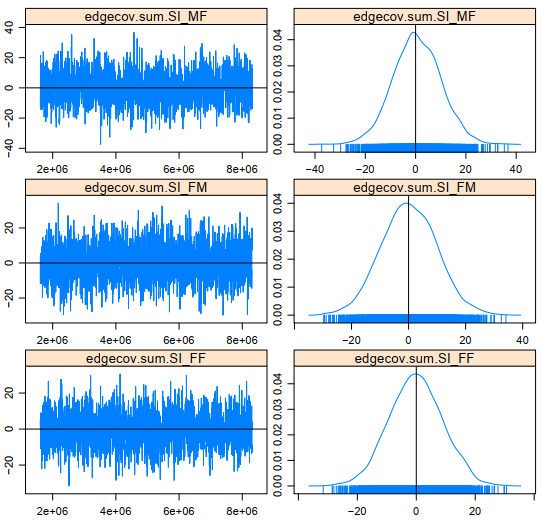

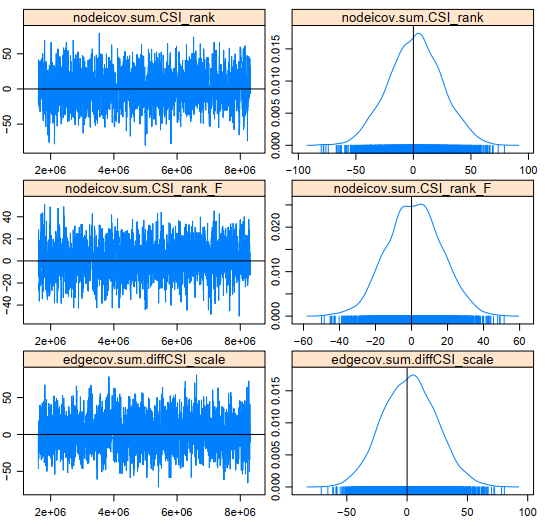

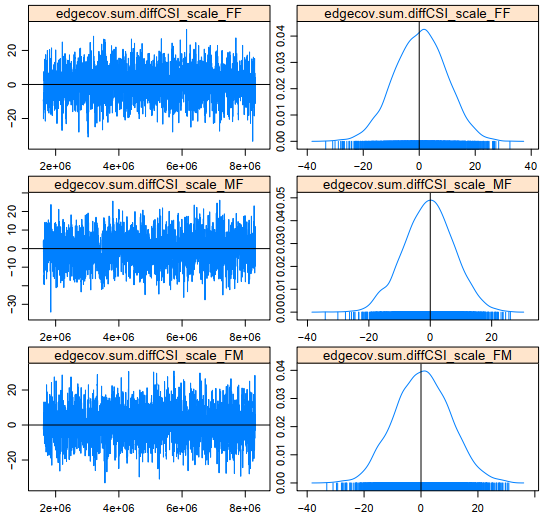

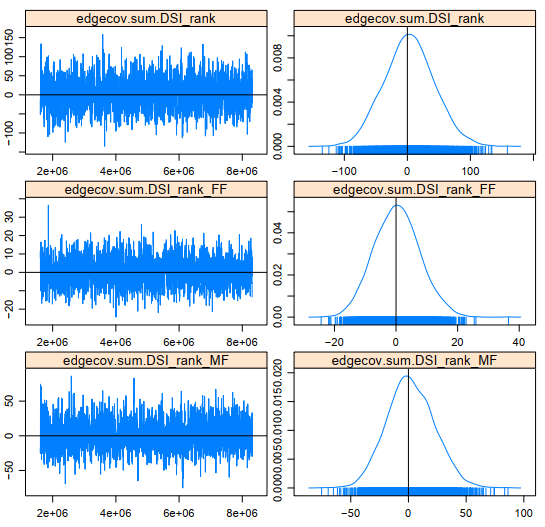

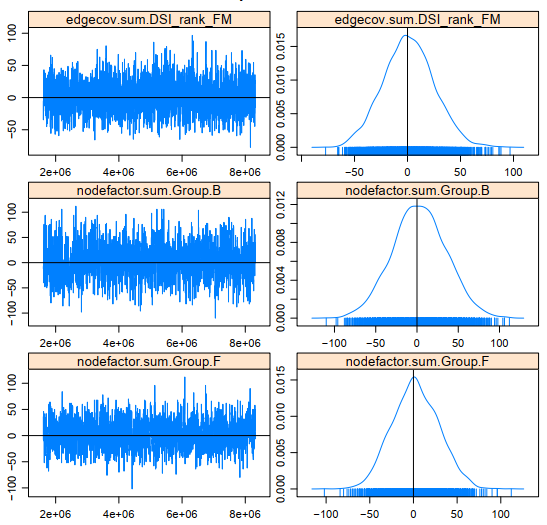

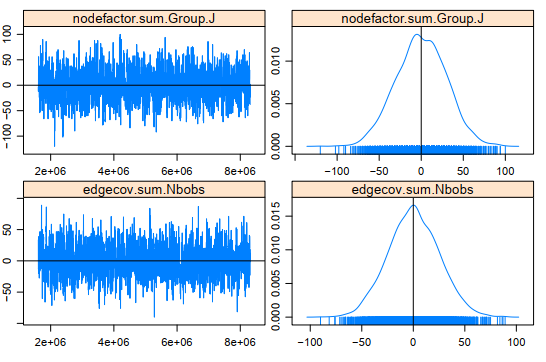
