## Supplementary File S1 for "Males are more sensitive to their audience than females when scent-marking in the redfronted lemur"

**Supplementary File S1:** Diagnostics for the 6 generalised linear mixed models

Supplementary File S1.A. Stability of the estimates for the male models when considering a) the 3m radius, b) the 5m radius, and c) the 10m radius.

a.

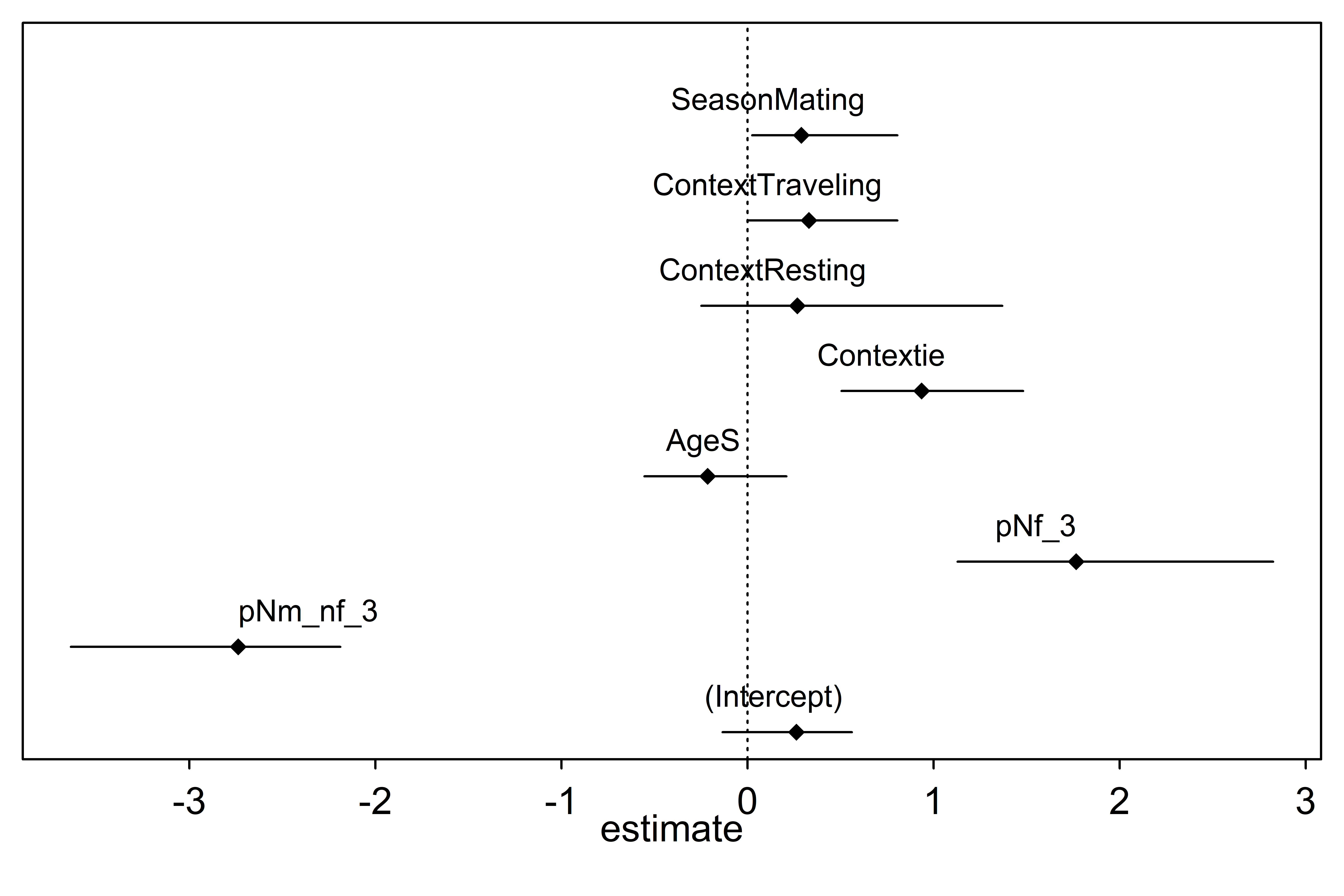

b.

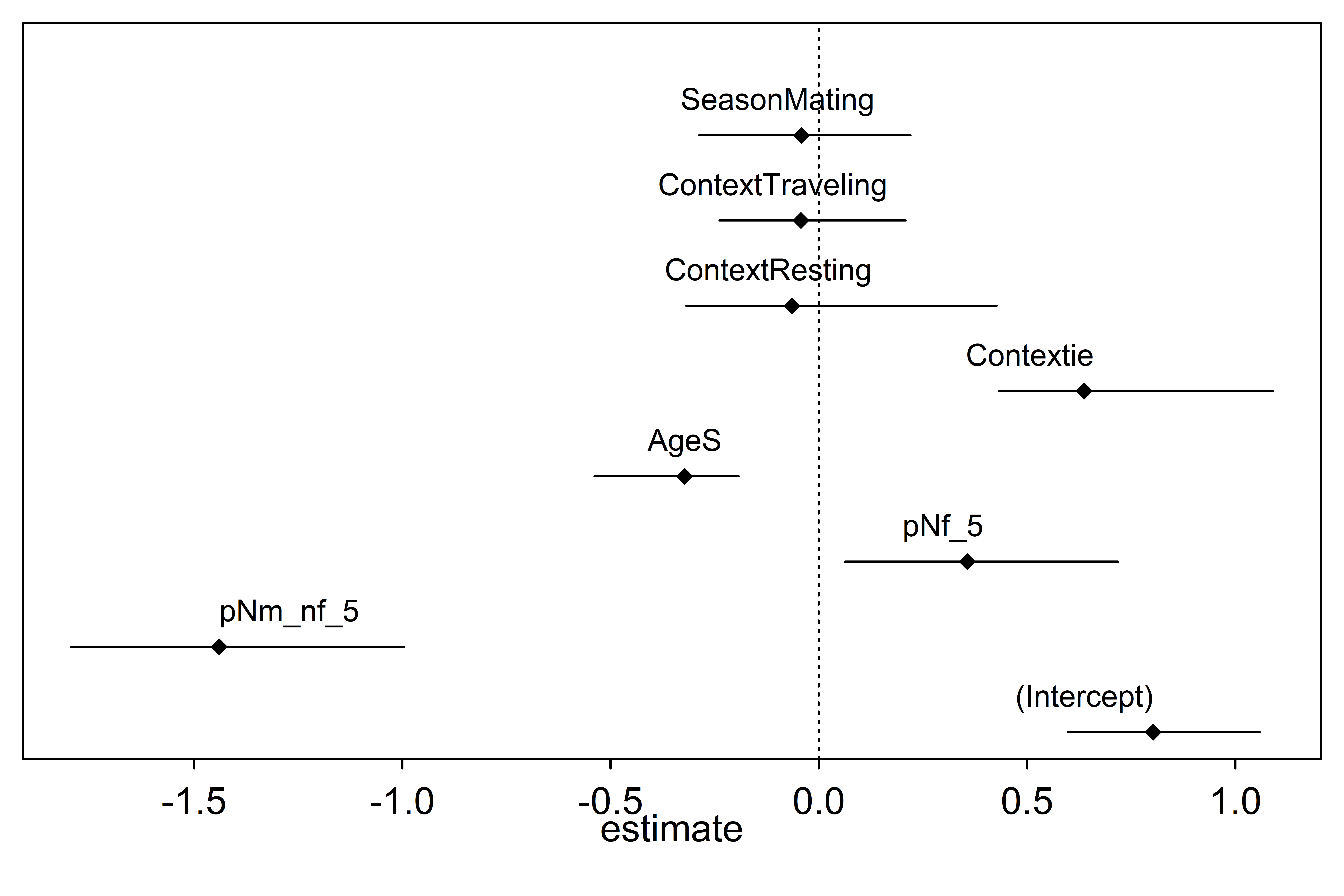

c.

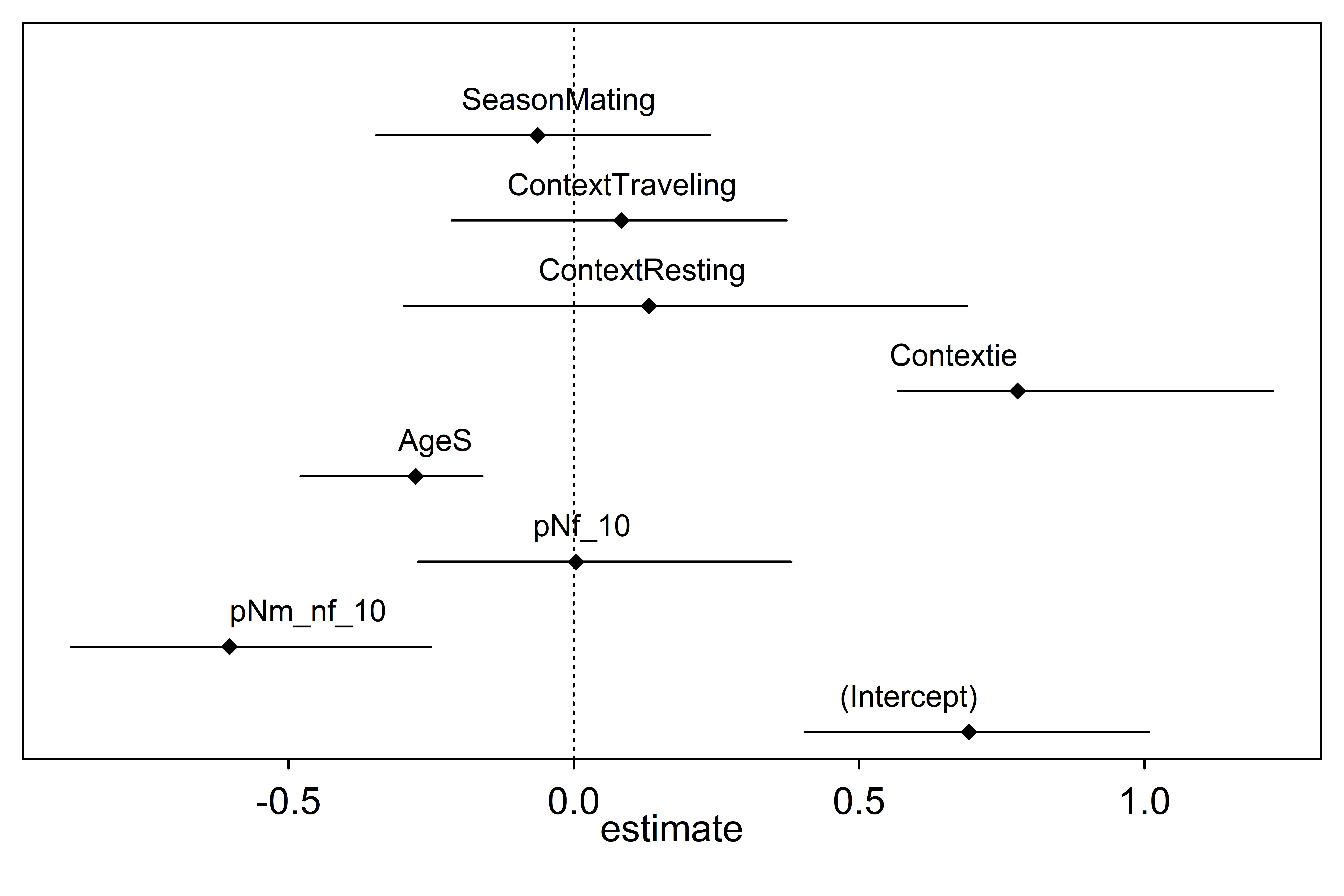

Supplementary File S1.B. Variation Inflation Factors for the male models

|  | 3m | 5m | 10m |
| --- | --- | --- | --- |
| Proportion of females | 1.35 | 1.54 | 1.76 |
| Proportion of males | 1.15 | 1.27 | 1.40 |
| Age | 1.04 | 1.03 | 1.04 |
| Context | 1.07 | 1.13 | 1.11 |
| Season | 1.13 | 1.09 | 1.04 |

Supplementary File S1.C. Stability of the estimates for the female models when considering a) the 3m radius, b) the 5m radius, and c) the 10m radius.

a.

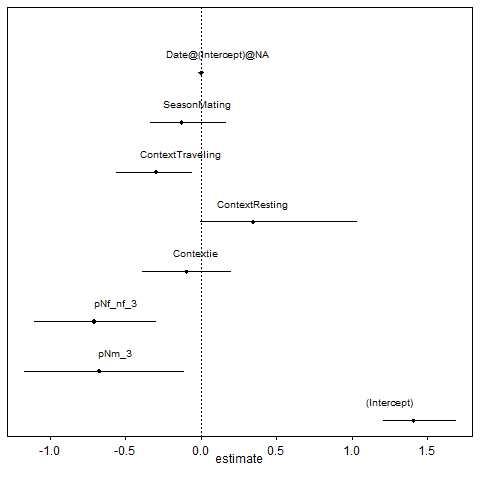

b.

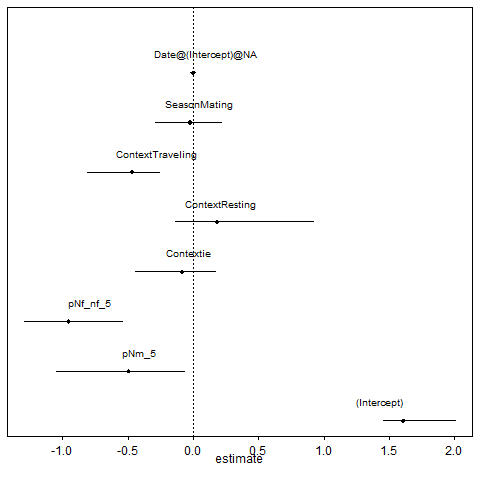

c.

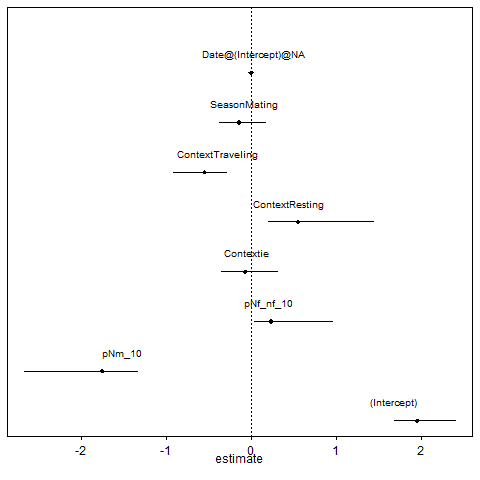

Supplementary File S1.D. Variation Inflation Factors for the female models

|  | 3m | 5m | 10m |
| --- | --- | --- | --- |
| Proportion of females | 2.12 | 2.20 | 1.80 |
| Proportion of males | 2.03 | 2.25 | 1.83 |
| Context | 1.07 | 1.09 | 1.11 |
| Season | 1.08 | 1.09 | 1.07 |
