## Supplementary File S2 for "Males are more sensitive to their audience than females when scent-marking in the redfronted lemur"

**Supplementary File S2: Bayesian analyses**

Because our models seem to suffer singularity issues, we further applied a Bayesian method as recommended by the authors of the “lme4” package ^1^. This approach should allow both regularising the model via informative priors and giving estimates and credible intervals for all parameters that average over the uncertainty in the random-effects parameters.

We fitted hierarchical linear models with the same structure as the ones described above using the R package “bmrs” ^2^ based on Stan modelling language ^3^. We used the default priors, namely a Student’s t-distribution (ν=3, μ=0, σ=10) for standard deviation for the likelihood function and unbiased priors for regression coefficients. Ten sampling chains were run for 6000 iterations and a warm-up period of 2000 iterations each. As the different traces overlap (Figure S2.A and Figure S2.B below) and that all Ȓ-values were equal to 1.00 we considered that the chains sufficiently converged. Additionally, to check whether these models reflect the observed data, we compared the samples from the posterior predictive distribution to the observed data using the function “pp_check()”. We reported for each variable the expected values under the posterior distribution and its 95% credible intervals (CIs). We judged that we had compelling evidence of an effect when 0 was not included in the 95% CI. Below are presented the tables and figure corresponding to table 2 (Table S2.A), table 3 (Table S2.B) and figure 1 (Figure S2.C) of the main manuscript but including the results of the Bayesian approach for comparison with the frequentist approach.

Table S2.A: Results of the models of the effects of audience composition within 3, 5 and 10m radius, age, context and season on the probability that a male scent-mark when passing a marking spot.

|  | **Frequentist approach** | | | | | | | | | | **Bayesian approach** | | | |
| --- | --- | --- | --- | --- | --- | --- | --- | --- | --- | --- | --- | --- | --- | --- |
|  | **audience radius** | **estimate** | **standard errors** | **lower CI** | **upper CI** | **chi-squared** | **df** | **p-value** | **minimum** | **maximum** | **estimate** | **Estimation error** | **Lower CI** | **Upper CI** |
| Intercept | 3m  5m  10m | 0.26  0.73  0.80 | 0.51  0.55  0.63 | -0.94  -0.58  -0.27 | 1.64  2.72  1.94 | - | - | - | -0.13  0.36  0.34 | 0.56  1.09  1.34 | 0.40  1.08  1.29 | 0.61  0.75  0.89 | -0.79  -0.34  -0.32 | 1.60  2.64  3.21 |
| **Proportion of males in the audience** | 3m  5m  10m | -2.74  -1.83  -0.83 | 1.02  0.84  0.64 | -7.65  -5.11  -1.88 | -0.83  -0.21  0.65 | 6.23  4.55  1.18 | 1  1  1 | **0.013**  **0.033**  0.277 | -3.64  -2.37  -1.34 | -2.19  -0.93  -0.37 | -3.22  -2.10  -0.76 | 1.34  1.26  1.05 | -5.99  -4.70  -2.86 | -0.71  0.34  1.35 |
| Proportion of females in the audience | 3m  5m  10m | 1.77  0.40  0.06 | 1.06  0.78  0.77 | -0.02  -1.60  -1.32 | 5.71  2.73  1.35 | 2.81  0.20  0.00 | 1  1  1 | 0.094  0.658  0.995 | 1.13  -0.06  -0.50 | 2.82  1.06  0.41 | 1.74  0.41  -0.34 | 1.27  1.03  1.03 | -0.54  -1.53  -2.44 | 4.50  2.56  1.64 |
| Age - Sub-adult | 3m  5m  10m | -0.21  0.21  -0.15 | 0.53  0.66  0.52 | -0.64  -1.18  -1.25 | 1.25  1.92  0.69 | 0.10  0.04  0.42 | 1  1  1 | 0.341  0.848  0.515 | -0.55  -0.38  -0.48 | 0.21  0.69  1.13 | -0.23  -0.30  -0.29 | 0.65  0.70  0.77 | -1.51  -1.65  -1.75 | 1.06  1.10  1.31 |
| Context - Inter-group encounter | 3m  5m  10m | 0.94  0.62  0.77 | 0.79  0.59  0.61 | -0.45  -0.81  -0.14 | 4.65  3.65  2.42 | 1.53  1.60  2.60 | 3  3  3 | 0.795  0.659  0.457 | 0.50  0.37  0.41 | 1.48  1.16  1.18 | 1.24  1.27  1.40 | 0.89  0.91  0.93 | -0.33  -0.32  -0.22 | 3.22  3.26  3.47 |
| Context - Resting | 3m  5m  10m | 0.27  0.13  0.10 | 0.88  1.72  1.14 | -2.43  -6.95  -1.28 | 4.85  11.01  1.95 |  |  |  | -0.25  -0.48  -0.52 | 1.37  5.54  5.96 | 0.36  -0.02  0.01 | 1.25  1.35  1.48 | -2.03  -2.61  -2.84 | 3.04  2.83  3.12 |
| Context - Traveling | 3m  5m  10m | 0.33  -0.09  -0.08 | 0.57  0.57  0.56 | -0.98  -1.86  -0.88 | 1.97  1.52  1.12 |  |  |  | 0.00  -0.34  -0.45 | 0.81  0.37  0.26 | 0.42  0.00  -0.09 | 0.65  0.74  0.76 | -0.83  -1.45  -1.64 | 1.75  1.47  1.35 |
| Season - Mating | 3m  5m  10m | 0.29  0.38  0.04 | 0.67  0.78  0.70 | 1.49  1.68  1.42 | 2.96  4.43  1.26 | 0.12  0.13  0.01 | 1  1  1 | 0.511  0.714  0.914 | 0.02  0.02  0.47 | 0.81  0.70  0.51 | 0.25  -0.07  -0.26 | 0.90  1.04  1.14 | -1.58  -2.17  -2.63 | 2.05  2.00  1.98 |

Table S2.B: Results of the models of the effects of audience composition within 3, 5 and 10m radius, age, context and season on the probability that a female scent-marked when passing a marking spot.

|  | **Frequentist approach** | | | | | | | | | | **Bayesian approach** | | | |
| --- | --- | --- | --- | --- | --- | --- | --- | --- | --- | --- | --- | --- | --- | --- |
|  | **audience radius** | **estimate** | **standard errors** | **lower CI** | **upper CI** | **chi-squared** | **df** | **p-value** | **minimum** | **maximum** | **estimate** | **Estimation error** | **Lower CI** | **Upper CI** |
| Intercept | 3m  5m  10m | 1.41  1.61  1.95 | 0.39  0.45  0.68 | 0.71  0.89  0.99 | 2.44  2.99  5.00 | - | - | - | -0.17  1.46  1.68 | 0.48  2.00  2.40 | 1.66  1.87  1.99 | 0.55  0.61  0.71 | 0.65  0.73  0.68 | 2.80  3.14  3.48 |
| Proportion of males in the audience | 3m  5m  10m | -0.67  -0.50  -1.75 | 0.84  0.85  1.00 | -2.62  -2.49  -5.45 | 1.34  1.35  -0.06 | 0.64  0.34  3.01 | 1  1  1 | 0.425  0.558  0.083 | -3.67  -1.05  -2.67 | -2.12  -0.07  -1.34 | -1.05  -0.73  -1.61 | 1.21  1.18  1.08 | -3.48  -3.03  -3.81 | 1.29  1.61  0.48 |
| Proportion of females in the audience | 3m  5m  10m | -0.71  -0.95  0.23 | 0.71  0.66  0.73 | -2.50  -2.60  -1.48 | 0.78  0.48  2.45 | 1.00  2.13  0.11 | 1  1  1 | 0.319  0.145  0.745 | 0.95  -1.29  0.03 | 2.79  -0.55  0.96 | -0.50  -0.84  0.22 | 0.95  0.89  0.78 | -2.33  -2.57  -1.26 | 1.43  0.93  1.81 |
| Context - Inter-group encounter | 3m  5m  10m | -0.10  -0.09  -0.07 | 0.53  0.53  0.56 | -1.20  -1.24  -1.73 | 1.23  1.27  2.05 | 1.01  1.30  1.71 | 3  3  3 | 0.799  0.730  0.634 | 0.11  -0.44  -0.35 | 0.76  0.17  0.30 | -0.02  -0.10  -0.10 | 0.79  0.78  0.80 | -1.51  -1.60  -1.61 | 1.62  1.48  1.55 |
| Context - Resting | 3m  5m  10m | 0.34  0.18  0.55 | 0.64  0.65  0.79 | -0.92  -1.30  -1.11 | 2.75  2.53  6.57 | 0.07  0.00  0.06 | 1  1  1 | 0.790  0.962  0.813 | 0.35  -0.14  0.21 | 1.36  0.92  1.44 | 0.42  0.30  0.70 | 0.93  0.95  0.97 | -1.36  -1.52  -1.13 | 2.31  2.28  2.71 |
| Context - Traveling | 3m  5m  10m | -0.30  -0.47  -0.55 | 0.49  0.51  0.61 | -1.44  -1.79  -2.65 | 0.80  0.64  0.91 |  |  |  | -0.37  -0.81  -0.92 | 1.45  -0.26  -0.29 | -0.40  -0.59  -0.53 | 0.64  0.67  0.70 | -1.69  -1.92  -1.92 | 0.85  0.72  0.86 |
| Season - Mating | 3m  5m  10m | -0.13  -0.02  -0.14 | 0.48  0.49  0.59 | -1.18  -1.08  -1.69 | 1.29  1.35  2.21 |  |  |  | -0.03  -0.29  -0.38 | 0.73  0.22  0.16 | -0.07  0.06  0.03 | 0.74  0.74  0.81 | -1.42  -1.31  -1.48 | 1.49  1.61  1.74 |

**Figure S2.A:** traces overlap for the male model a) in the 3m radius, b) in the 5m radius and c) in the 10m radius.

a)	b)	c)

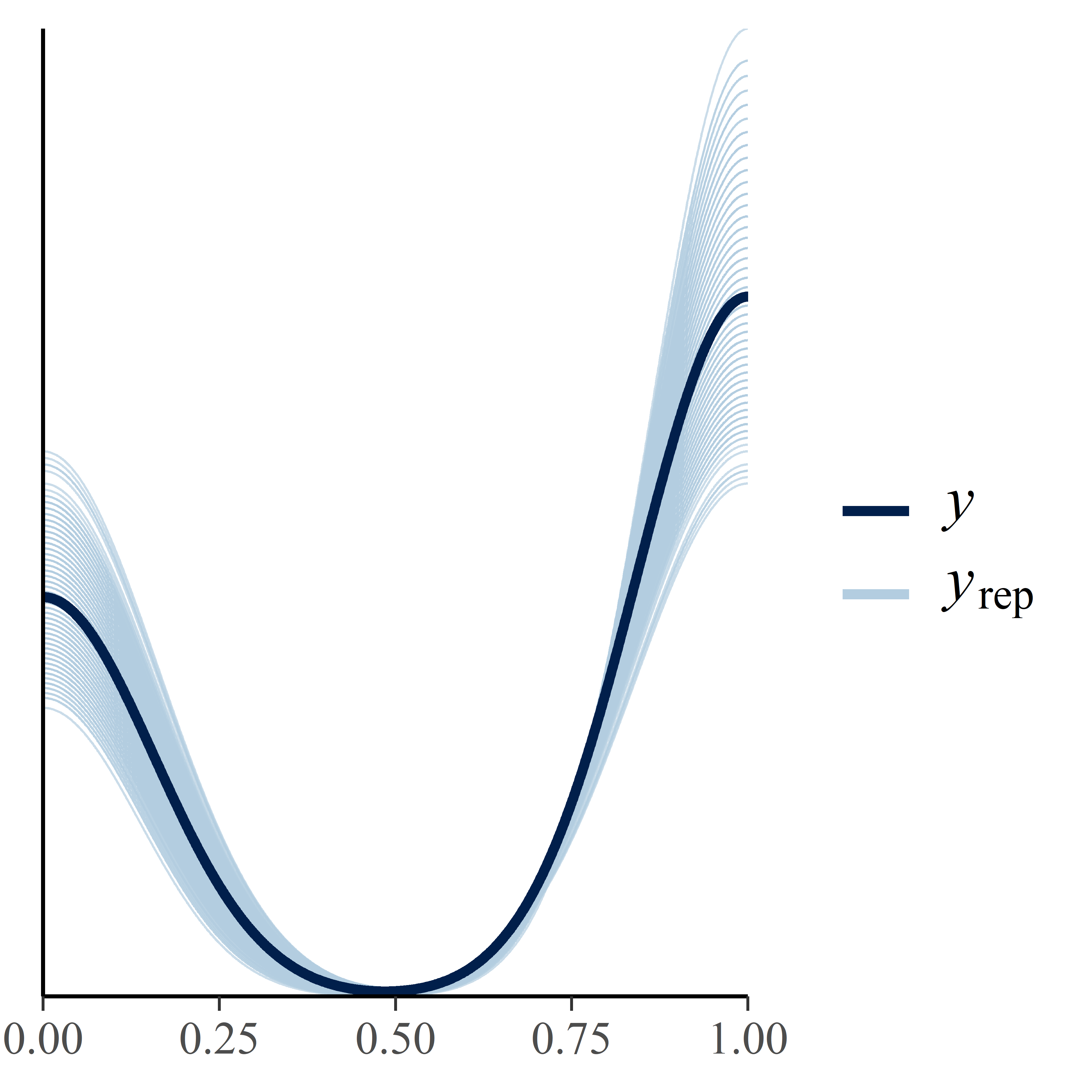

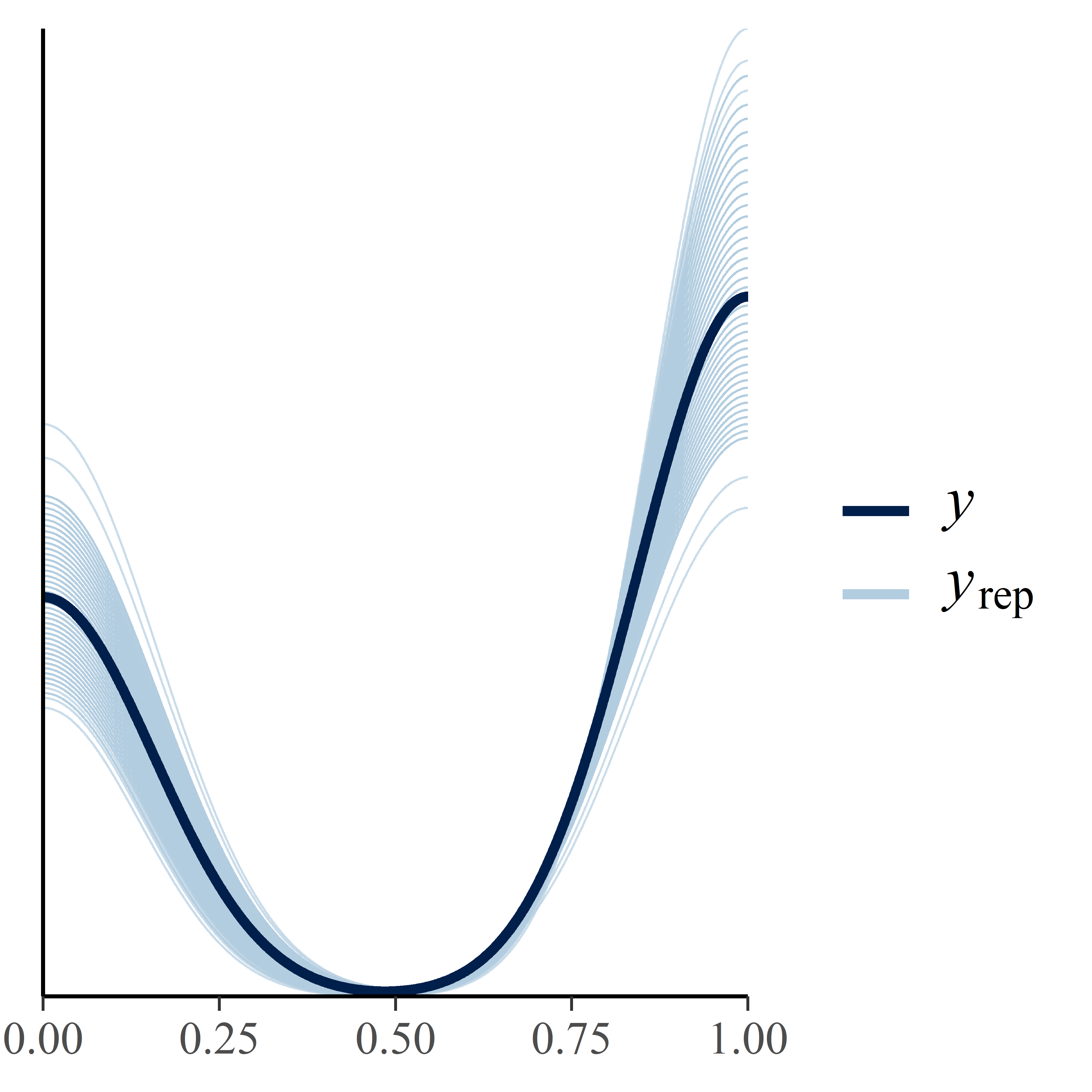

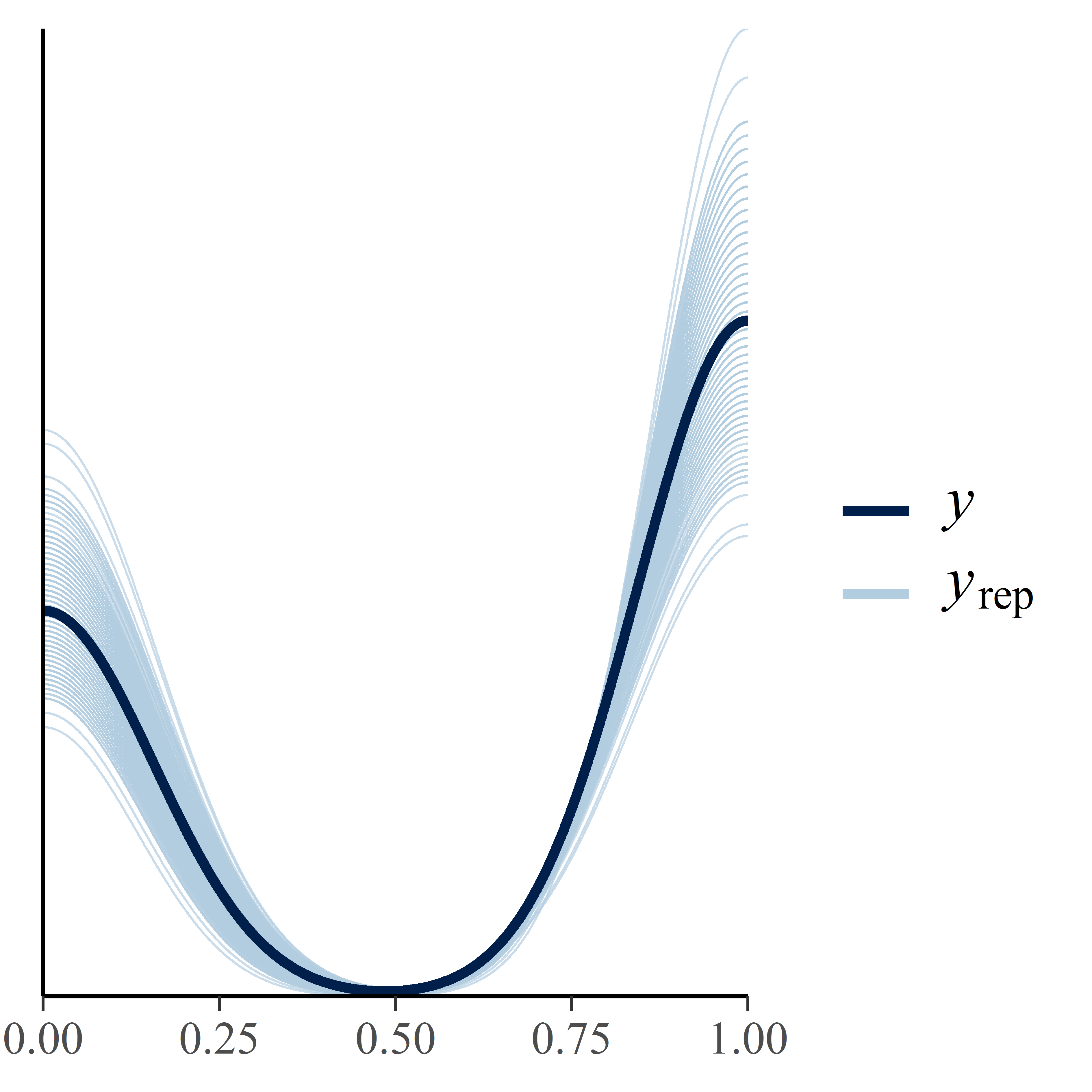

**Figure S2.B:** traces overlap for the female model a) in the 3m radius, b) in the 5m radius and c) in the 10m radius.

a)	b)	c)

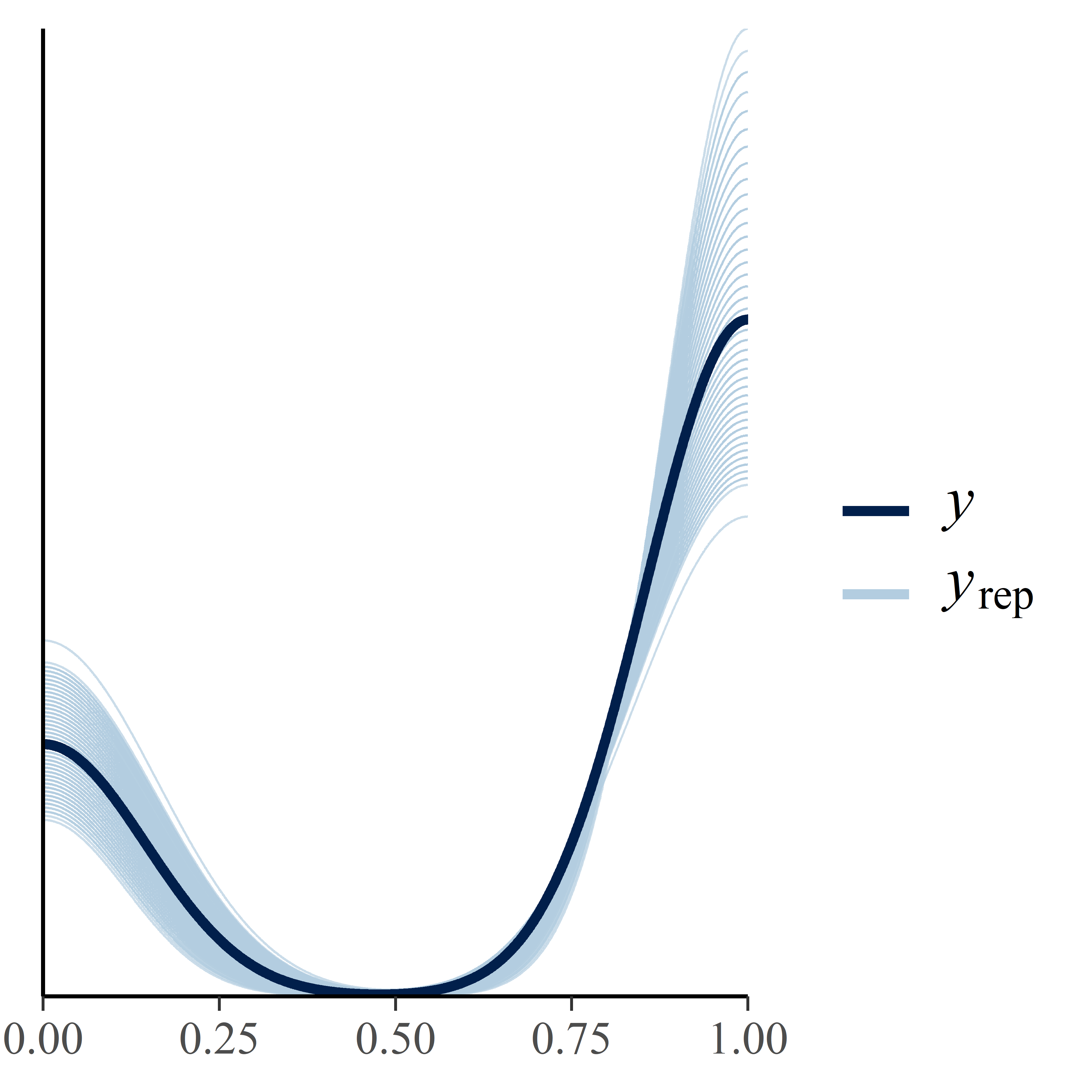

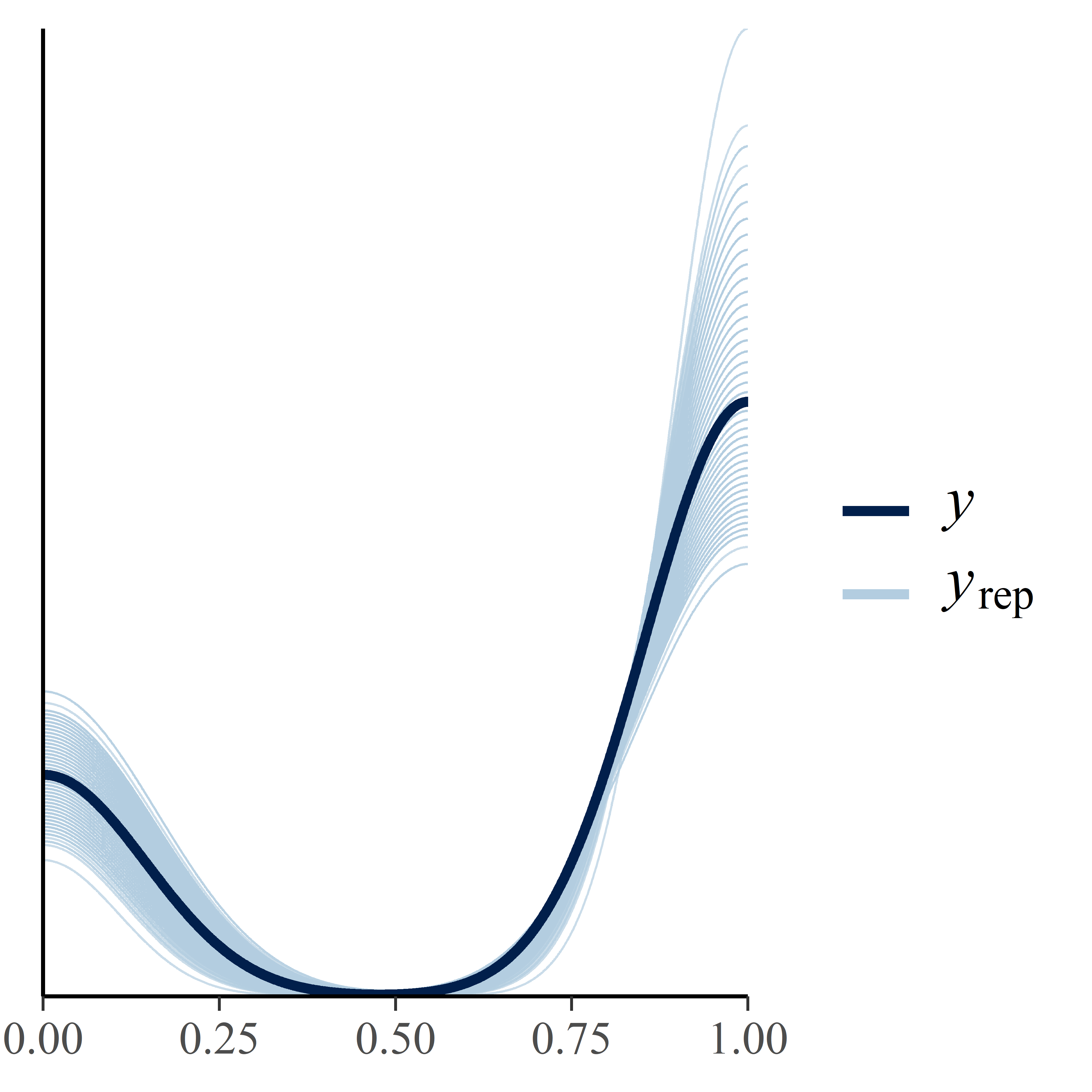

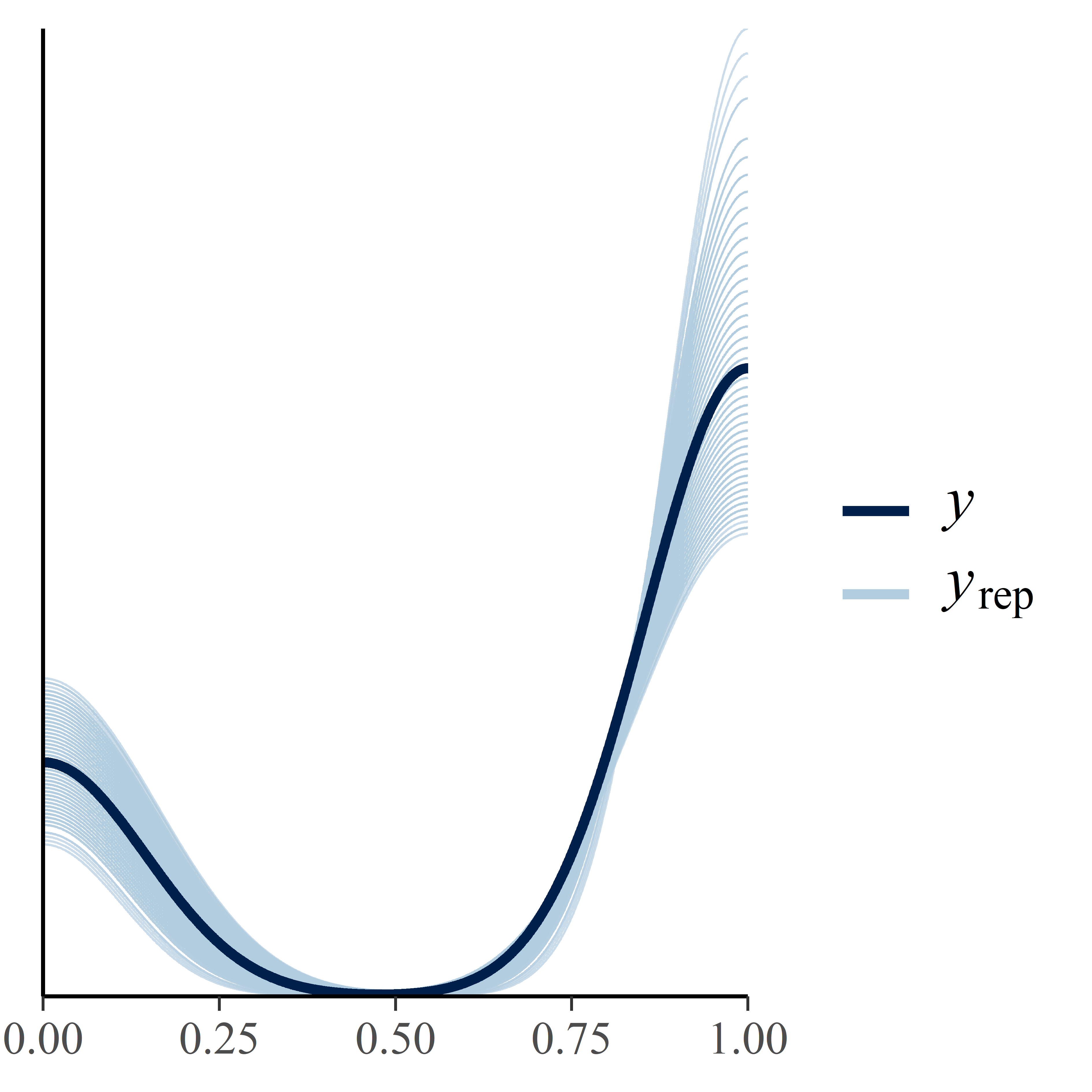

**Figure S2.C:** Probability that a male genital-mark depending on the proportion of males present in a)3m, b)5m, c)10m. Colours correspond to the different individuals (n=14) and the size of the circle corresponds to the number of observations (n=165). Below each graph, we present the corresponding expected values under the posterior distribution and its 95% credible intervals (CIs) obtained with the Bayesian approach.

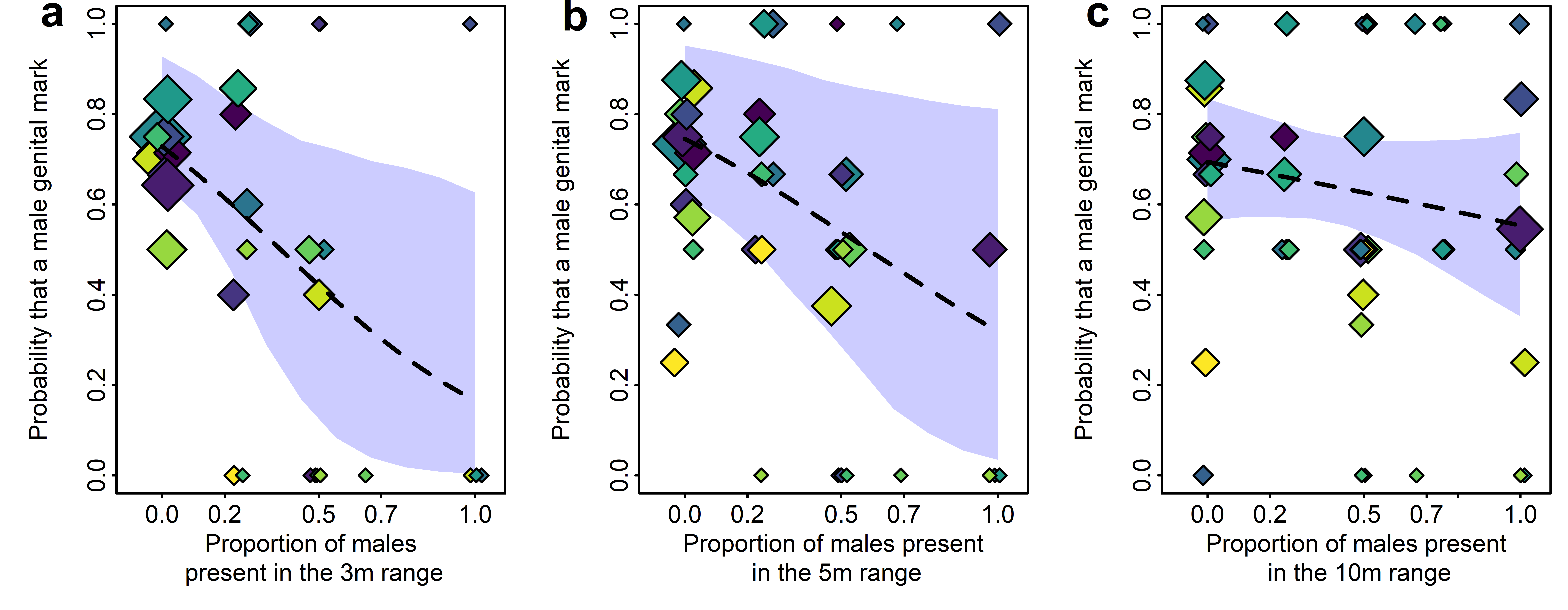

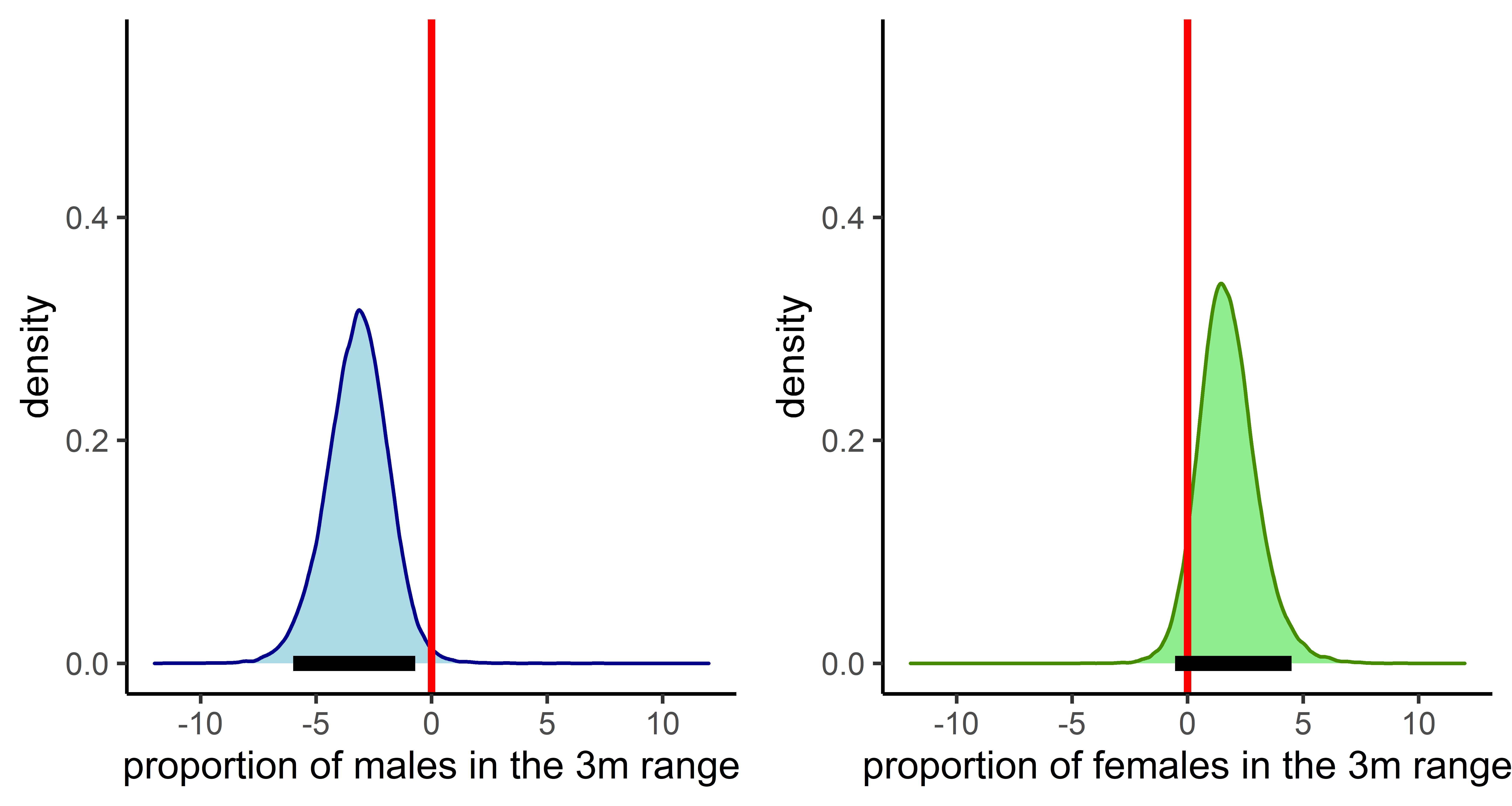

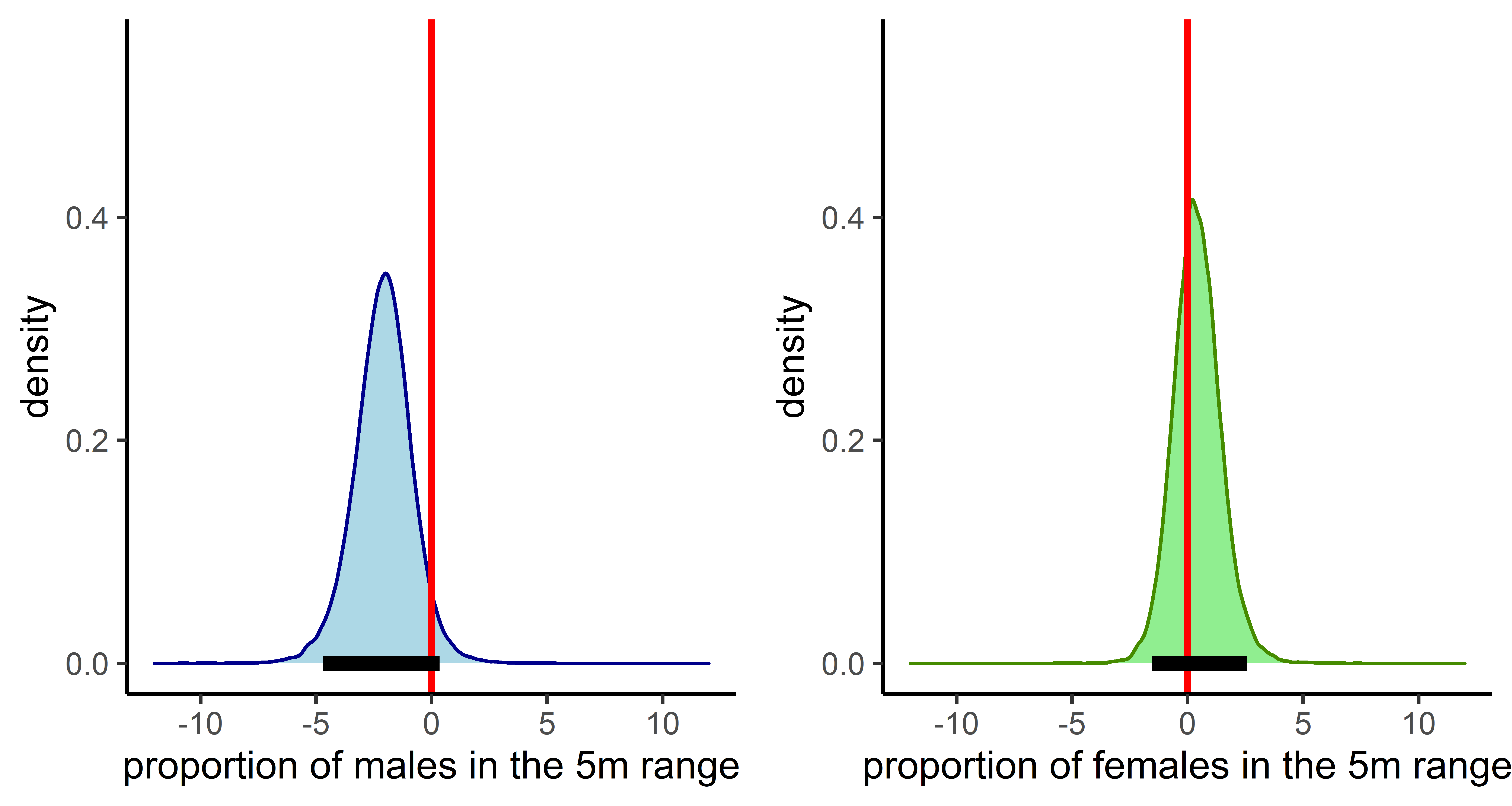

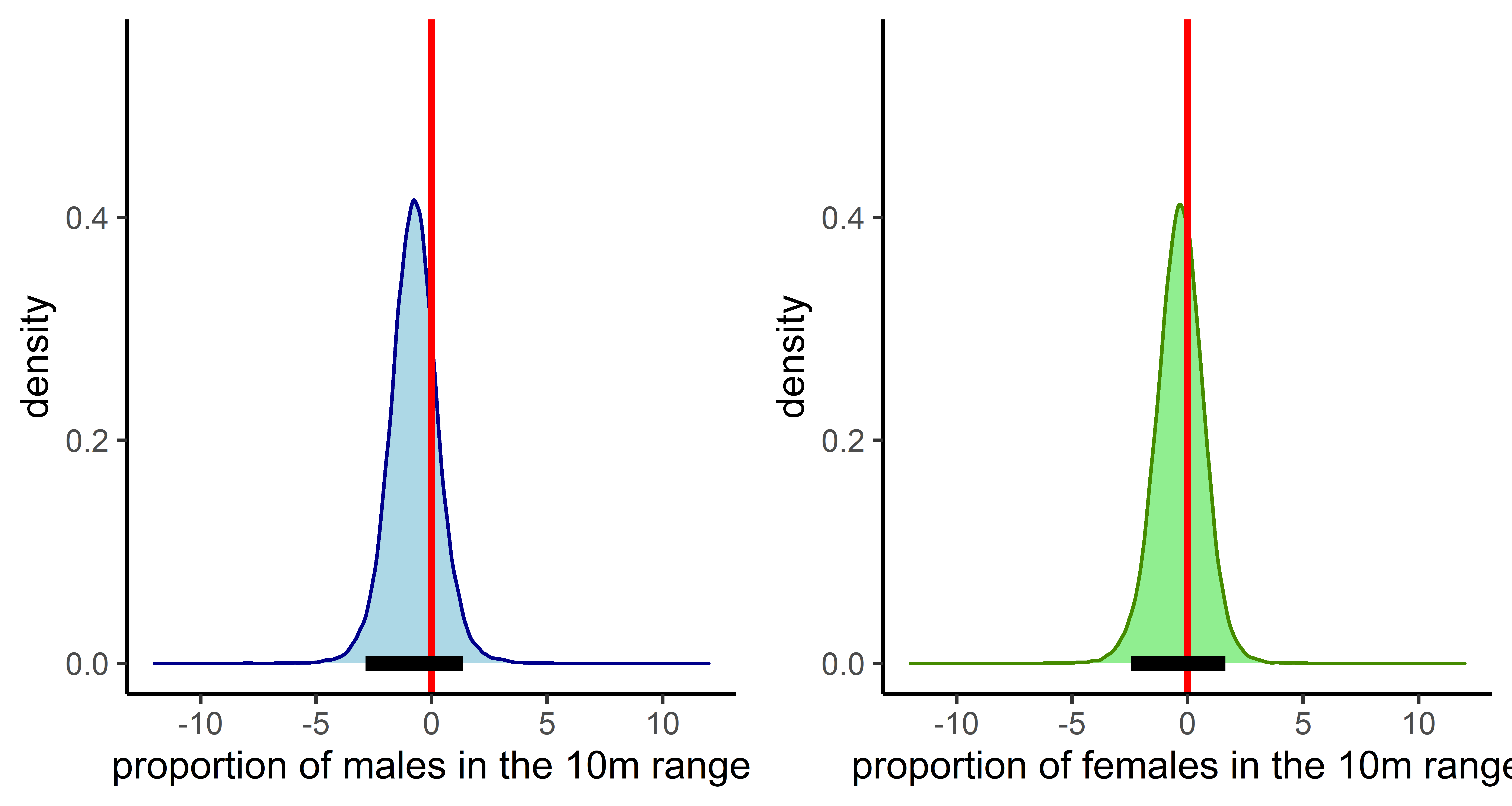
